# Collagen remodeling dictates pancreatic cancer bioenergetics and outcome through DDR1 activation or degradation

**DOI:** 10.1101/2022.04.02.486837

**Authors:** Hua Su, Fei Yang, Rao Fu, Brittney Trinh, Nina Sun, Junlai Liu, Avi Kumar, Jacopo Baglieri, Jeremy Siruno, Stephen Dozier, Ajay Nair, Aveline Filliol, Sara Brin Rosenthal, Jennifer Santini, Christian M. Metallo, Anthony Molina, Robert F. Schwabe, Andrew M. Lowy, David Brenner, Beicheng Sun, Michael Karin

## Abstract

Pancreatic ductal adenocarcinoma (PDAC) is a highly desmoplastic, aggressive cancer that frequently progresses by liver metastasis^1^. Cancer-associated fibroblasts (CAF), extracellular matrix (ECM), and type I collagen (Col I) support^2–5^ or restrain PDAC progression and may impede blood supply and nutrient availability^6–8^. The dichotomous role of the stroma in PDAC, and the mechanisms through which it influences patient survival and enables desmoplastic cancers escape nutrient limitation remain poorly understood. Here we show that matrix metalloprotease (MMP)-cleaved or intact Col I (cCol I and iCol I, respectively) exert opposing effects on PDAC bioenergetics, macropinocytosis (MP), tumor growth and liver metastasis. While cCol I activates DDR1 (discoidin domain receptor-1)-NF-κB-p62-NRF2 signaling to promote PDAC growth, iCol I triggers DDR1 degradation and restrains PDAC growth. Patients whose tumors are enriched in iCol I and low in DDR1 and NRF2 have improved median survival compared to those enriched in cCol I, DDR1 and NRF2. Inhibition of DDR1-stimulated NF-κB or mitochondrial biogenesis blocked tumorigenesis in wildtype mice but not in mice expressing MMP-resistant Col I. In summary, the diverse effects of tumor stroma on PDAC growth, metastasis, and patient survival are mediated through the Col I-DDR1-NF-κB-NRF2-mitochondrial biogenesis pathway, presenting multiple new opportunities for PDAC therapy.

## Main

Retrospective clinical studies suggest that PDAC patients with a fibrogenic but inert tumor stroma, defined by extensive ECM deposition, low expression of the myofibroblast marker α-SMA and low MMP activity, have improved progression-free survival (PFS) as compared to patients whose tumors are populated by fibrolytic stroma, defined by low collagen fiber content, high α-SMA expression and MMP activity^9^. How the stromal state affects clinical outcome is unknown. Moreover, previous investigations of stromal influence on PDAC growth and progression yielded conflicting results, assigning stroma and CAFs as either tumor supportive^5, 10^ or restrictive^7, 8^. It is likely that the failure of stromal-targeted PDAC therapies^11, 12^ is due, at least in part, to unrecognized pathways that result in tumor-promoting or tumor-suppressive stromal subgroups, thus requiring precision medicine rather than one-size-fits-all approaches.

## Cleaved and intact Col I differentially affect PDAC growth

To further investigate how the fibrolytic stroma may affect PDAC outcome, we compared survival between patients with high and low fibrolysis, using a signature of 6 collagen cleaving MMPs (MMP1, 2, 8, 9, 13, 14) as a surrogate for collagenolysis. This analysis showed that high MMP expression correlated with poor survival (Extended Data Fig.1a). One main target for MMPs in desmoplastic tumors is Col I, the prevalent ECM protein. We also used antibodies that distinguish iCol I from cCol I (¾ Col I) (Extended Data Fig. 1b) to stratify a cohort of 106 resected PDAC patients (see below) and correlate the Col I state of their tumors with survival. Again, the results pointed to Col I remodeling as a strong prognostic factor as patients whose tumors were enriched in cCol I had poorer median survival (Fig. 1a). To understand the basis for these results and mimic an inert tumor stroma, low in cCol I, we transplanted mouse PDAC cells into mice with either wildtype *Col1α1^+/+^* (Col I^WT^), or MMP-resistant Col I generated by two amino acids (AA) substitutions in the α1 subunit that block Col I cleavage by MMPs^13^, *Col1α1^r/r^* (Col I^r/r^). Col I^r/r^ mice develop more extensive hepatic fibrosis (HF) than Col I^WT^ mice, but despite the hepatocellular carcinoma (HCC) supportive functions of HF^14, 15^, they poorly accommodate HCC growth due to unknown mechanisms^16^. Col I^WT^ and Col I^r/r^ mice were either orthotopically or intrasplenically (i.s.; as a model of liver metastasis) transplanted with mouse PDAC KPC960 (KPC) or KC6141 (KC) cells. Col I^r/r^ mice poorly supported primary pancreatic tumor growth or hepatic metastases, even though their pancreata were more fibrotic than Col I^WT^ pancreata. This finding persisted in mice pretreated with the pancreatitis inducer caerulein (CAE), which stimulated liver metastasis in Col I^WT^ pancreata (Fig. 1b, c, Extended Data Fig. 1c-e). Following i.s. transplantation, KPC or KC tumors in Col I^WT^ livers were larger in CCl_4_ pretreated mice, while tumor numbers and size were diminished in Col I^r/r^ livers, regardless of CCl_4_ pretreatment (Fig.1d, Extended Data Fig. 1f). As expected, Col I^r/r^ livers were more fibrotic than Col I^WT^ livers regardless of CCl_4_ pretreatment (Extended Data Fig. 1g). Primary PDAC and liver metastases were confirmed by staining with ductal (CK19), progenitor (SOX9) or proliferation (Ki67) markers (Extended Data Fig. 1d, e, h). Enhanced tumor growth in CAE or CCl_4_ pretreated Col I^WT^ mice suggested that tumor suppression in Col I^r/r^ mice is not simply due to space limitation imposed by excessive Col I buildup.

**Fig. 1:**
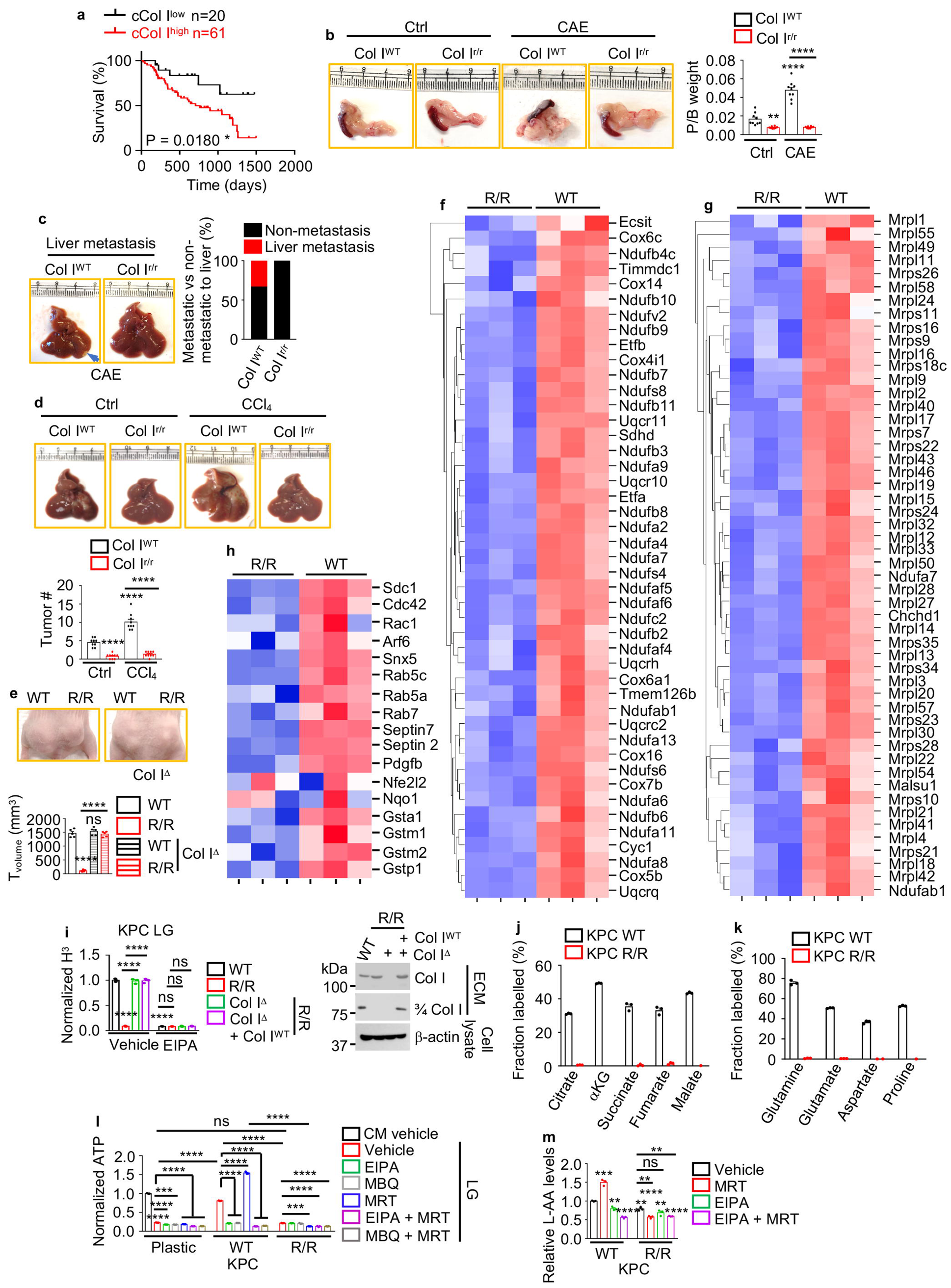
Col I state controls PDAC growth and metabolism. **a**, Overall survival of resected PDAC patients stratified according to cCol I expression determined by immunohistochemical (IHC) analysis in Fig. 3a. Significance was determined by log rank test. *p < 0.05. **b**, Pancreas morphology and weight relative to body weight (P/B) 4 weeks (wk) after orthotopic KPC cell transplantation into Col I^WT^ or Col I^r/r^ mice pretreated -/+ CAE. **c**, Liver morphology in above CAE-treated mice. Liver metastases were detected in 33% of CAE-treated Col I^WT^ mice. **d**, Liver gross morphology and tumor numbers (#) 2 wk after i.s. KPC cell transplantation into Col I^WT^ or Col I^r/r^ mice pretreated -/+ CCl_4_. **e**, Representative images, and sizes of s.c. tumors formed by human 1305 cells co-transplanted with WT, R/R, or Col I^Δ^ WT or R/R fibroblasts into *Nu/Nu* mice. **f**-**h**, Differentially expressed genes between KPC cells grown on WT or R/R ECM in low glucose (LG, 0.5 mM) for 24 h. Vertical rows: different specimens; horizontal rows: individual genes, colored according to log-transformed z-score units. TPM (transcripts per million). Blue: replicates with low expression (z-score = −2); red: replicates with high expression (z-score = +2). Mitochondrial electron transfer chain (ETC) genes (**f**), mitochondrial ribosome subunit genes (**g**), MP-related and NRF2-target genes (**h**). **i**, KPC cells grown on ^3^H-proline-labeled WT, R/R, Col I^Δ^ R/R or Col I^Δ^ R/R + Col I^WT^ ECM were incubated in LG -/+ EIPA (10.5 μM) for 24 h. ^3^H uptake was measured by liquid scintillation, normalized to cell number, and presented as ^3^H CPM relative to vehicle treated WT ECM plated KPC cells. Immunoblot (IB) analysis of iCol I and 3/4 Col I in the extracellular matrix (ECM) produced by indicated fibroblasts. (**j, k**) Fractional labeling (mole percent enrichment) of TCA cycle intermediates (**j**) and intracellular AA (**k**) in KPC cells incubated for 24 h in LG medium after plating on WT or R/R ECM deposited by fibroblasts labeled with [U-^13^C] glutamine for 5 days. (**l**) KPC cells plated on WT or R/R ECM, or plastic were incubated in LG -/+ EIPA, MRT68921 (MRT, 600 nM), MBQ-167 (MBQ, 500 nM), EIPA + MRT, MBQ + MRT for 24 h. Total cellular ATP was measured, normalized to cell number and presented relative to untreated plastic-plated cells. (**m**) Total AA in KPC cells plated on WT or R/R ECM and incubated in LG -/+ EIPA, MRT, EIPA + MRT for 24 h. Data normalized to cell number are presented relative to untreated WT ECM-plated cells. Results in **b** (n=8-9), **d** (n=9), **e** (n=5), (**i**, **l**, and **m**) (n=3 independent experiments), (**j**) and (**k**) (n=3 per condition) are mean ± SEM. Statistical significance was determined by a two-tailed t test. **p < 0.01, ***p < 0.001, ****p < 0.0001.

To determine the effect of Col I remodeling on human PDAC growth, we subcutaneously (s.c.) co-transplanted WT and R/R fibroblasts with a patient derived xenograft (PDX) cell line (1305) into immunocompromised *Nu/Nu* mice. WT fibroblasts enhanced tumor growth, whereas R/R fibroblasts inhibited tumor growth but lost their inhibitory activity after *Col1α1* gene ablation (Fig. 1e). *Col1α1* ablation did not affect the stimulatory activity of WT fibroblasts, suggesting a specific inhibitory function of noncleaved Col I.

## The Col I state controls PDAC metabolism

To determine the basis for blunted tumorigenesis in Col I^r/r^ mice, we plated KPC cells on ECM deposited by WT and R/R fibroblasts, incubated them in low glucose (LG) medium (to model nutrient restriction) and conducted RNA-sequencing (RNA-seq). Bioinformatic analysis revealed striking differences between cells cultured on WT vs. R/R ECM, with the former showing upregulation of sulfur AA metabolism, mammary gland morphogenesis, telomere maintenance and RNA processing mRNAs and the latter exhibiting upregulation of innate immunity and inflammation related mRNAs (Extended Data Fig. 2a). The most striking differences were in MP-related genes and nuclear and mitochondrial genes encoding mitochondrial ribosome subunits and electron transfer chain (ETC) components, which were upregulated by WT and suppressed by R/R ECM (Fig. 1f-h). Consistent with upregulation of MP genes by WT ECM, IKKα-deficient KC cells, which have high MP activity^17^, formed more tumors than parental KC cells in Col I^WT^ livers, while growing as poorly as parental KC cells in Col I^r/r^ livers (Extended Data Fig.1f).

**Fig. 2:**
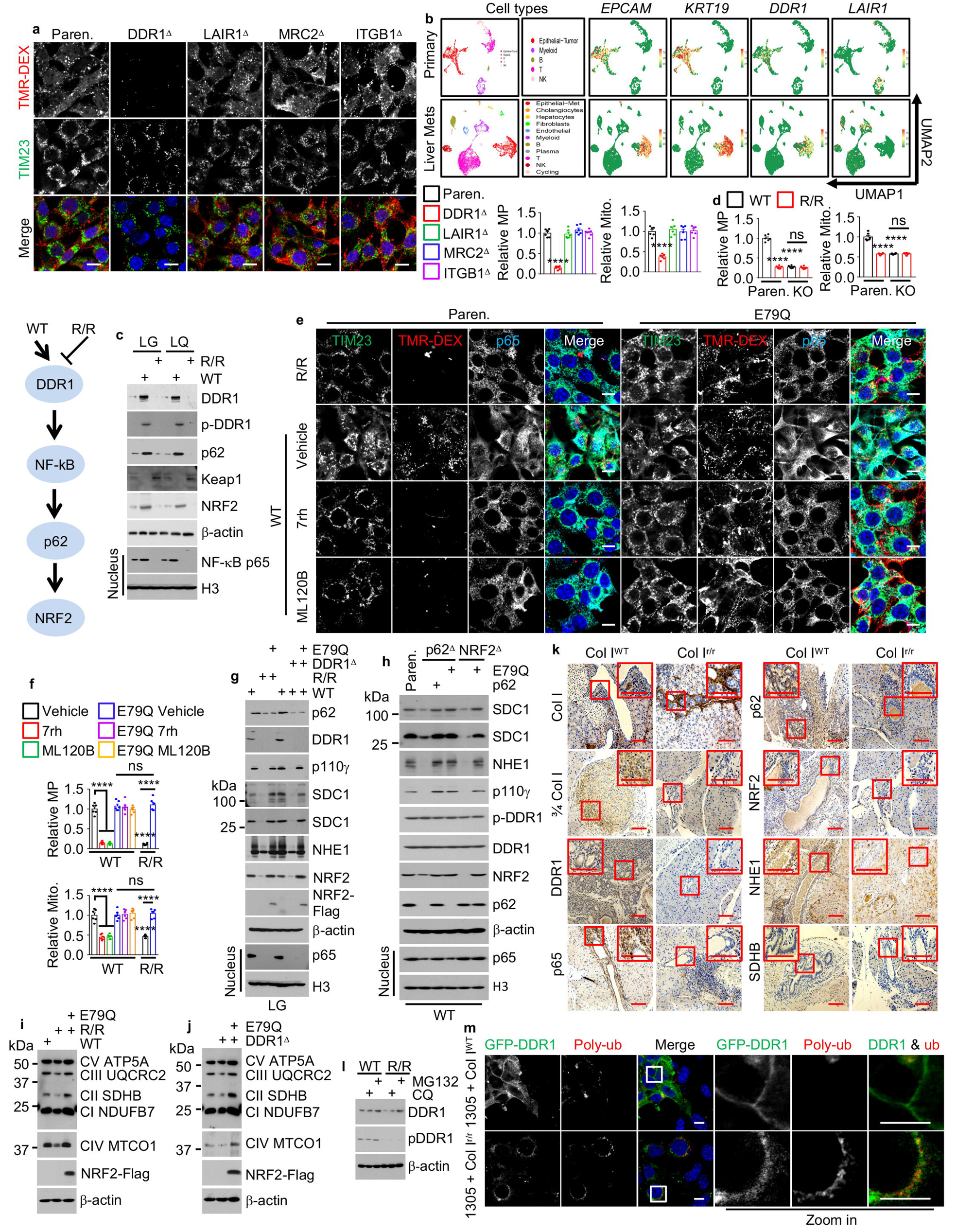
Col I controls MP and mitochondrial biogenesis through the DDR1-NRF2 axis. **a**, Representative images and quantification of mitochondria (Mito., TIM23) and MP in TMR-dextran (TMR-DEX, red)-incubated parental (Paren.) and variant KPC cells plated on WT ECM. **b**, UMAPs showing scRNA-seq data from 5 patients with primary PDAC (upper row) and 1 patient with PDAC liver metastasis (lower row), displaying the identified cell types and expression of *EPCAM*, *KRT19*, *DDR1* and *LAIR1* mRNAs. **c**, IB analysis of indicated proteins in KPC cells plated on plastic, WT or R/R ECM and incubated in LG or low glutamine (LQ, 0.2 mM) media for 24 h. The effects of WT and R/R ECM on DDR1 signaling are summarized on the left. **d**, Quantification of Mito. and MP in TMR-DEX-incubated Paren. and NRF2^Δ^ (KO) KPC cells plated on WT or R/R ECM and incubated in LG for 24 h. **e, f**, Representative images (**e**) and quantification (**f**) of Mito. and MP in TMR-DEX-incubated Paren. and NRF2^E79Q^ KPC cells plated on WT or R/R ECM and incubated in LG -/+ 7rh (500 nM), ML120B (10 μM) for 24 h. **g**, IB analysis of indicated proteins in Paren., NRF2^E79Q^, DDR1^Δ^, NRF2^E79Q^/DDR1^Δ^ KPC cells treated as (**d**). **h**, IB analysis of indicated KPC cells -/+ exogenous p62 or NRF2^E79Q^ transfections plated on WT ECM and incubated in LG for 24 h. **i**, IB analysis of ETC complexes I-V (CI-CV) in Paren. and NRF2^E79Q^ KPC cells treated as (**d**). **j**, IB analysis of ETC proteins in Paren., DDR1^Δ^, or NRF2^E79Q^/DDR1^Δ^ KPC cells plated on WT ECM and incubated in LG for 24 h. **k**, Representative IHC analysis of the indicated proteins in Col I^WT^ and Col I^r/r^ pancreata 4 wk after orthotopic transplantation of KPC cells. Boxed areas were further magnified. Scale bars, 100 μm. **l**, IB analysis of indicated proteins in KPC cells plated on WT or R/R ECM and incubated in LG -/+MG132 (10 μM) or chloroquine (CQ, 50 μM) for 24 h. **m**, Representative images showing colocalization of GFP-DDR1 and poly-ubiquitin (poly-ub) in GFP-DDR1 expressing 1305 cells cocultured with WT or R/R fibroblasts in LG for 24 h. Boxed areas were further magnified. Results in (**a**, **d**, and **f**) (n=6) are mean ± SEM. Statistical significance was determined by a two-tailed t test. ****p < 0.0001. (**a**, **e**, **m**), scale bars, 10 μm.

To assess effects of Col I on PDAC metabolism, we labeled WT and R/R fibroblasts with [^3^H] proline or [U-^13^C] glutamine for 5 days, during which the cells coated the plates with Col I-containing ECM. After decellularization, KPC or KC cells and variants thereof were plated and cultured for 24 h in LG. [^3^H] uptake in WT ECM plated cells was MP-dependent, indicated by sensitivity to MP inhibitors (NHE1i EIPA, PI3Kγi IPI549, CDC42/RACi MBQ-167) and NHE1 or SDC1 knockdowns (KD), and enhancement by the ULK1i MRT68921 (MRT)^17^. In contrast, ^3^H uptake in cells plated on R/R ECM was barely detectable and unaffected by MP inhibition (Fig. 1i, Extended Data Fig. 2b-d). Notably, *Col1α1* gene ablation or overexpression of cleavable Col I in ECM laying R/R fibroblasts restored ^3^H uptake (Fig. 1i). Strikingly, cells cultured on ^13^C-glutamine-labelled WT ECM took up glutamine and metabolized it, but cells plated on ^13^C-glutamine-labelled R/R ECM exhibited minimal glutamine uptake and metabolism (Fig. 1j, k). Congruently, WT ECM cultured cells had higher ATP and AA content, that was further increased by MRT treatment and reduced by MP blockade, as compared to R/R ECM cultured cells, whose low ATP and AA contents were barely affected by the MP inhibitors (Fig. 1l, m, Extended Data Fig. 2e-h). Col I ablation or WT Col I overexpression prevented the decline in ATP and AA (Extended Data Fig. 2f, h), suggesting that cCol I is an important signaling molecule that stimulates PDAC metabolism and energy generation.

## The cleaved to intact Col I ratio controls DDR1 to NRF2 signaling

KPC or human MIA PaCa-2 cells plated on WT ECM or co-cultured with WT fibroblasts in LG or low glutamine (LQ) exhibited high MP rates measured by tetramethylrhodamine-labeled high-molecular-mass dextran (TMR-DEX) uptake, whereas cells plated on R/R ECM or co-cultured with R/R fibroblasts exhibited low MP rates (Extended Data Fig. 3a, b). Furthermore, culturing KPC cells on WT ECM for 24 h. markedly upregulated MP-related proteins and NRF2 relative to culture on plastic, but culturing with R/R ECM had the opposite effect (Extended Data Fig. 3c). Similar differences in NRF2 and MP-related proteins, mRNAs and MP activity were displayed by KPC tumors growing in Col I^WT^ or Col I^r/r^ livers or pancreata (Extended Data Fig. 3d-f). Mitochondria are important for cancer growth by generating energy for macromolecular synthesis^18^. Consistent with the RNA-seq data, mitochondria and ETC proteins were decreased in PDAC cells grown on R/R ECM or in Col I^r/r^ pancreata (Extended Data Fig. 3g-i).

**Fig. 3:**
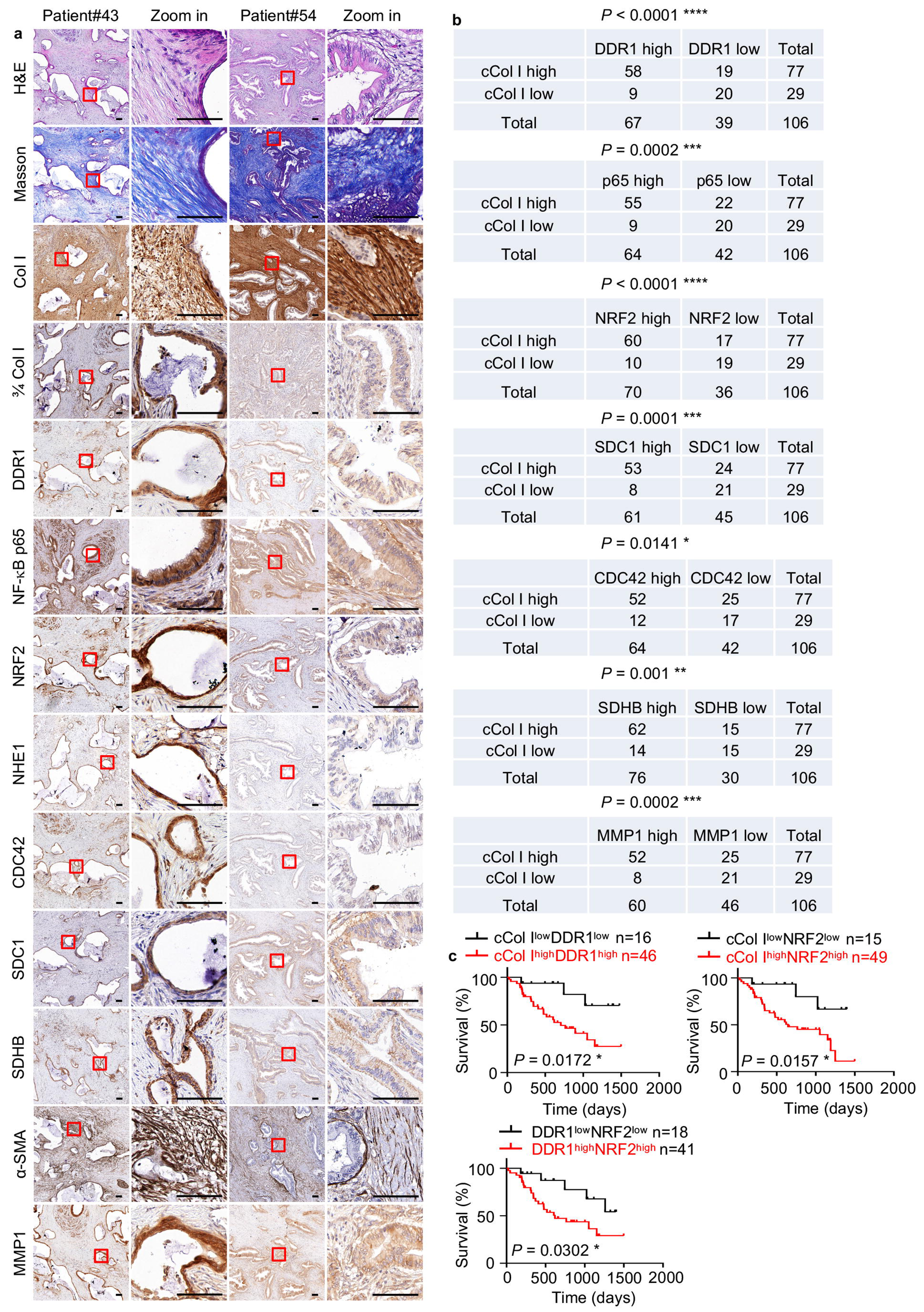
Elevated cCol I and increased DDR1 and NRF2 predict poor patient survival. **a**, Representative IHC analysis of resected human PDAC tissues. The boxed areas were further magnified. Scale bars, 100 μm. **b**, Correlations between cCol I (3/4 Col I) and indicated proteins in the above human PDAC tissue array were analyzed by a Chi-square test. *p < 0.05, **p < 0.01, ***p < 0.001, ****p < 0.0001. **c**, Comparisons of overall survival between patients stratified according to cCol I, DDR1 and NRF2 expression. Significance was determined by log rank test. *p < 0.05.

The human PDAC stroma consists of both intact and cleaved collagens. To recapitulate this setting and determine how the iCol I to cCol I balance affects PDAC metabolism, we mixed R/R with WT (R:W) or Col I^Δ^ R/R (R:KO) fibroblasts to generate ECM with different iCol I and cCol I amounts, confirmed with isoform specific antibodies. KPC cells were plated on the ECM preparations, kept in LG for 24 h, and their MP rates, mitochondria numbers and nuclear NRF2 were evaluated. Nondegradable Col I at 6:4 (R:W) or 4:6 (R:KO) ratios and above inhibited MP and reduced mitochondria numbers and nuclear NRF2 (Extended Data Fig. 3j, k). This experiment suggests that iCol I inhibits MP and mitochondria biogenesis, which are stimulated by different cleaved collagens, not only cCol I.

To investigate how the Col I state regulates MP and mitochondrial biogenesis, we systematically ablated (Extended Data Fig. 4a) all known collagen receptors expressed by KPC cells, MRC2, DDR1, LAIR-1, and β1 integrin^19^. The only receptor whose ablation inhibited MP activity and mitochondrial biogenesis (Fig. 2a) was DDR1, a collagen activated receptor tyrosine kinase (RTK)^20^, which single cell RNA-seq analysis showed to be highly expressed in primary and liver metastatic human PDAC cells, marked by *EPCAM* and *KRT19* mRNA expression (Fig. 2b, Extended Data Fig. 4b). Other collagen receptor mRNAs were either not expressed in PDAC (*LAIR1*, *MRC2*) or had a broader distribution (*ITGB1*). Whereas WT ECM stimulated DDR1 expression and phosphorylation, R/R ECM dramatically downregulated DDR1 and its downstream effector NF-κB^21^ as well as p62 (Fig. 2c), encoded by an NF-κB target^22^. The inhibitory effect of iCol I was probably missed in previous studies of DDR1 activation and signaling, which used artificially fragmented acid solubilized collagens as DDR1 ligands^23^. Consistent with p62 induction, WT ECM decreased KEAP1 and upregulated NRF2, whereas R/R collagen had the opposite effects (Fig. 2c). We wondered whether Col I affects MP and mitochondrial biogenesis through the DDR1-NF-κB-p62-NRF2 cascade. Indeed, R/R ECM, NRF2, DDR1or IKKβ inhibition decreased MP activity, 3/4 Col I fragment uptake, NRF2 nuclear localization, mitochondria number, and expression of MP-related and mitochondrial ETC proteins (Fig. 2d-j, Extended Data Fig. 4c-j). Overexpression of an activated NRF2^E79Q^ variant in KPC cells reversed the inhibitory effects of R/R ECM, DDR1i or IKKβi but did not restore/affect DDR1 expression/phosphorylation and p65 nuclear localization (Fig. 2e-j, Extended Data Fig. 4f-j). Consistent with these data, pancreatic and liver tumors from Col I^r/r^ mice showed more extensive iCol I expression but no cCol I and lower DDR1, p65, p62, NRF2, NHE1 and SDHB (mitochondrial marker) compared to tumors from Col I^WT^ mice (Fig. 2k, Extended Data Fig. 5a). These results suggest that Col I controls MP and mitochondrial biogenesis through the DDR1-NF-κB-p62-NRF2 cascade. As myofibroblast-specific Col I ablation enhances intrahepatic PDAC growth^7^, we examined how ECM from Col I^Δ^ fibroblasts affects MP and DDR1 signaling. We found that Col I^Δ^ ECM behaved just like WT ECM, stimulating MP, mitochondrial biogenesis and DDR1 phosphorylation, effects that were blocked by DDR1 ablation (Extended Data Fig. 5b-d). These results fit earlier descriptions of DDR1 as a general collagen receptor^20^, with other collagens in Col I^Δ^ fibroblasts acting as ligands.

**Fig. 4:**
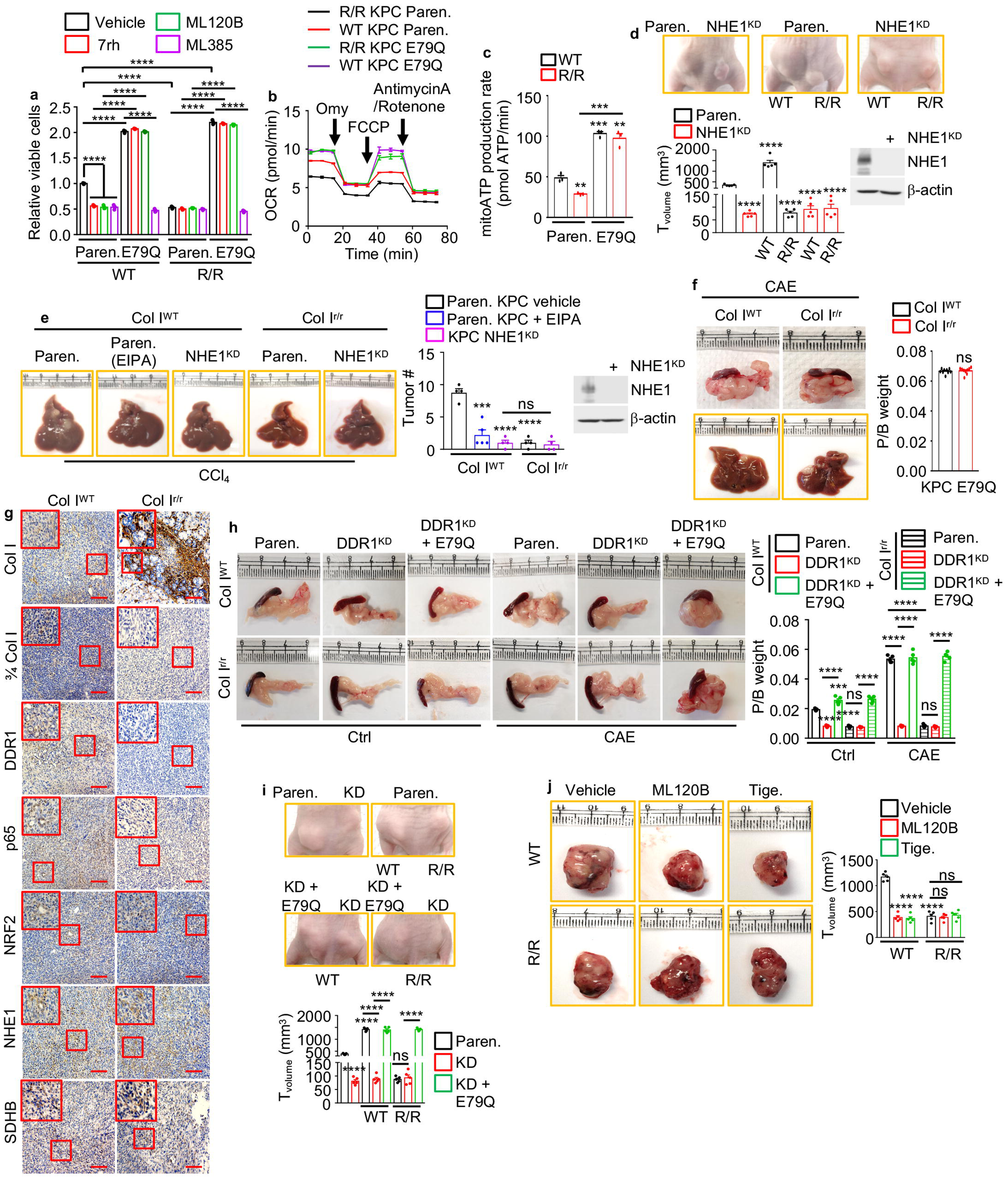
The therapeutically targetable DDR1-NF-κB-NRF2 axis controls PDAC growth and metabolism. **a**, Paren. and NRF2^E79Q^ KPC cells were plated on WT or R/R ECM and incubated in LG medium -/+ 7rh, ML120B, ML385 (10 μM). Total viable cells were measured as above, and data are presented relative to the untreated Paren. cells on WT ECM. **b**, Oxygen-consumption rates (OCR) of Paren. or NRF2^E79Q^ KPC cells plated on WT or R/R ECM and incubated in LG for 24 h before and after oligomycin (Omy), FCCP, and rotenone/antimycin A additions. **c**, Mito. ATP production calculated from the curves in (**b**). **d**, Representative images and sizes of Paren. and NHE1^KD^ MIA-PaCa-2 tumors s.c. grown -/+ WT or R/R fibroblasts in nude mice. IB analysis of NHE1 in MIA PaCa-2 cells is shown. **e**, Liver morphology and tumor numbers (#) 2 wk after i.s. transplantation of Paren. or NHE1^KD^ KPC cells into CCl_4_-pretreated Col I^WT^ and Col I^r/r^ mice -/+ EIPA treatment. IB analysis of NHE1 in KPC cells is shown on the right. **f**, Liver and pancreas morphologies and weight 4 wk after orthotopic transplantation of KPC NRF2^E79Q^ cells into CAE-pretreated Col I^WT^ and Col I^r/r^ mice. **g,** IHC analysis of pancreatic sections from above mice. Boxed areas were further magnified. Scale bars, 100 μm**. h,** Pancreas morphology and weight 4 wk after orthotopic transplantation of indicated KPC cells into Col I^WT^ and Col I^r/r^ mice pretreated -/+ CAE. **i**, Representative images and sizes of s.c. tumors generated by Paren., DDR1^KD^ (KD), NRF2^E79Q^/KD 1305 cells -/+ WT or R/R fibroblasts in nude mice. **j**, Representative images and sizes of s.c. MIA-PaCa tumors grown with WT or R/R fibroblasts in nude mice -/+ ML120B or tigecycline (Tige.) treatments. Results in (**a**) (n=3 independent experiments), (**c**) (n=3 per condition), (**d**, **h**-**j**) (n=5), (**e**) (n=4-5), and (**f**) (n=9) are mean ± SEM. Statistical significance was determined by a two-tailed t test. **p < 0.01, ***p < 0.001, ****p < 0.0001.

## Intact Col I triggers DDR1 proteasomal degradation

DDR1 expression and function vary in different cancer stages and types^24–29^. *Ddr1* mRNA was increased by culture on R/R ECM (Extended Data Fig. 5j), implying that diminished DDR1 protein expression in these cultures is post-transcriptional. Indeed, MG132, a proteasome inhibitor, but not the lysosomal inhibitor chloroquine, rescued DDR1 expression but not autophosphorylation (Fig. 2l). Notably, GFP-DDR1 showed strong cell surface localization and little polyubiquitin colocalization in 1305 cells cocultured with WT fibroblasts but was cytoplasmic and strongly colocalized with polyubiquitin in R/R fibroblast cocultures (Fig. 2m). Unlike DDR1 in triple negative breast cancer (TNBC)^28^, no shedding of the DDR1 extracellular domain was detected (Extended Data Fig. 5e). Our results therefore reveal a new mode of DDR1 regulation in PDAC and probably other desmoplastic cancers.

## NRF2 controls mitochondrial biogenesis and respiration

ECM from fibroblasts treated with the FDA-approved MMPi Ilomastat behaved like R/R ECM (Extended Data Fig. 5f, g), indicating that the results were not unique to the Col I^R^ variant. R/R ECM also decreased mitochondria in autophagy-deficient PDAC cells (Extended Data Fig. 5h), indicating reduced mitochondrial content is not due to mitophagy. In addition, colocalization of mitochondria and polyubiquitin which marks mitophagy was rarely observed (Extended Data Fig. 5i). Expression of TFAM, a key activator of mtDNA transcription, replication and biogenesis^30^ was downregulated in R/R ECM cultured PDAC cells, but *Nrf1* (unrelated to NRF2) mRNA, PGC1α protein and AMPK activity, which also stimulate mitochondrial biogenesis^31^, were upregulated (Extended Data Fig. 5j, k). The latter results match the low ATP content of R/R ECM cultured cells. *In silico* analysis revealed putative NRF2 binding sites in the *Tfam* promoter region to which NRF2 was recruited in cells plated on WT ECM or on NRF2^E79Q^ expression (Extended Data Fig. 5l, m). These results suggest that NRF2 is critical for cCol I stimulated MP and mitochondrial biogenesis.

## Elevated intact Col I and reduced DDR1 and NRF2 correlate with improved survival

Immunohistochemical (IHC) analysis of surgically resected human PDAC showed that most tumors (77/106) contained high amounts of 3/4 Col I and most of these tumors exhibited higher DDR1 (58/77), NF-κB p65 (55/77), NRF2 (60/77), SDC1 (53/77), CDC42 (52/77), SDHB (62/77), α-SMA (56/77) and MMP1 (52/77) staining than cCol I^low^ tumors (Fig. 3a, b, Extended Data Fig. 6a, b), suggesting that PDACs with fibrolytic stroma have high MP activity and mitochondrial content than tumors with inert stroma. Moreover, DDR1 and p65, DDR1 and NRF2, p65 and NRF2, NRF2 and MP proteins (NHE1, SDC1 or CDC42) or NRF2 and SDHB showed strong positive correlations (Extended Data Fig. 6b), suggesting that the fibrolytic stroma stimulates MP and mitochondrial biogenesis through the DDR1-NF-κB-NRF2 cascade also in human PDAC. Importantly, patients with cCol I^high^ and DDR1^high^, cCol I^high^ and NRF2^high^ or DDR1^high^ and NRF2^high^ tumors had considerably worse median survival than patients with low expression of these markers. These results are consistent with observations made in our preclinical PDAC models, suggesting that the fibrolytic stroma may drive human PDAC recurrence via NRF2-mediated MP and mitochondrial biogenesis.

## Therapeutic targeting of the DDR1-NF-κB-NRF2-tumor metabolism cascade

Increasing iCol I in the ECM inhibited cellular DNA synthesis (Extended Data Fig. 7a). Parental, NRF2^E79Q^, or IKKα^KD^ PDAC cells were plated on WT or R/R ECM, incubated in LG, and treated with DDR1(7rh), IKKβ (ML120B), NRF2 (ML385) or MP (NHE1^KD^ or EIPA, IPI549, MBQ-167) inhibitors. While WT ECM increased and R/R ECM decreased parental PDAC cell growth, inhibition of MP, DDR1, IKKβ or NRF2 decreased growth on WT ECM (Fig. 4a, Extended Data Fig. 7b-f). NRF2^E79Q^ expressing cells grew faster than parental cells and were resistant to R/R ECM, DDR1i or IKKβi but not NRF2i. IKKα^KD^ cells with high MP rates and nuclear NRF2 also grew faster than the parental cells on WT ECM but were more sensitive to R/R ECM and MP inhibitors (Extended Data Fig. 7b, c). MP, DDR1, IKKβ or NRF2 inhibition did not decrease low parental cell growth on R/R ECM (Fig. 4a, Extended Data Fig. 7b-f). Moreover, parental KPC or 1305 cells plated on WT ECM were more sensitive to the mitochondrial protein synthesis inhibitor tigecycline than cells on R/R ECM or DDR1^KD^ cells grown on WT ECM (Extended Data Fig. 7g). NRF2^E79Q^ cells showed higher oxygen consumption (OCR) and mitochondrial ATP production rates than parental cells, which were diminished by R/R ECM but only in the parental cells (Fig. 4b, c). Thus, the fibrolytic stroma may support PDAC cell growth through Col I stimulated MP and mitochondrial biogenesis. *In vivo*, R/R fibroblasts inhibited human PDAC (MIA PaCa-2) tumor growth, but WT fibroblasts were stimulatory. NHE1 ablation or EIPA inhibited tumor growth with or without co-transplanted WT fibroblasts or in WT livers but had little effect on tumors growing with R/R fibroblasts or in Col I^r/r^ livers (Fig. 4d, e). Tumors growing with WT fibroblasts were more fibrotic than tumors without added fibroblasts and small tumors growing with R/R fibroblasts had the highest collagen content (Extended Data Fig. 8a), indicating that Col I deposition enhances PDAC growth only when it is MMP cleaved. NRF2^E79Q^ cells in Col I^r/r^ hosts exhibited similar growth, NRF2, NHE1 and SDHB expression and liver metastases as cells growing in Col I^WT^ hosts, despite low DDR1 and p65 expression (Fig. 4f, g, Extended Data Fig. 8b).

In TNBC, DDR1 supports progression by aligning collagen fibers to elicit immune exclusion^28^. Using second harmonic generation (SHG) measurements, we observed no change in collagen fiber alignment between tumors from Col I^WT^ and Col I^r/r^ pancreata or parental and DDR1^KD^ tumors (Extended Data Fig. 8c). Accordingly, DDR1 ablation inhibited tumor growth, p65, p62, NRF2, NHE1 and SDHB expression in WT pancreas but did not reduce it further in Col I^r/r^ pancreas (Fig. 4h, Extended Data Fig. 8d). NRF2^E79Q^ rescued tumor growth and NHE1 and SDHB, but not p65 or p62, expression in DDR1^KD^ cells regardless of Col I status. Similar results were observed in immunodeficient mice (Fig. 4i), indicating that the effects of Col I-DDR1 interaction differ between PDAC and TNBC. Importantly, inhibition of IKKβ or mitochondrial protein synthesis decreased growth of tumors cotransplanted with WT, but not R/R, fibroblasts (Fig. 4j), illustrating two other ways of targeting PDAC with fibrolytic stroma.

## Discussion

This study shows that Col I remodeling is prognostic for PDAC patient survival. In preclinical models, Col I remodeling modulated tumor growth and metabolism via a DDR1-NF-κB-p62-NRF2 cascade, activated by cCol I and inhibited by iCol I. DDR1 activation by collagens and downstream NF-κB activation were described^21, 23^. However, it was previously unknown that iCol I triggers DDR1 polyubiquitylation and proteasomal degradation, indicating that DDR1 distinguishes cleaved from intact collagens, the latter capable of restraining tumor metabolism and growth. Although DDR1i inhibit mouse PDAC growth^32^, DDR1’s ability to control tumor metabolism by stimulating MP and mitochondrial biogenesis was unknown. It is perplexing, however, that DDR1, a rather weak RTK^20, 33^, exerts such a strong effect on metabolism in PDAC cells that express more potent RTKs, such as EGFR and MET. Perhaps this is due to the stronger NF-κB activating capacity of DDR1 relative to other RTKs. Indeed, IKKβ inhibition was as effective as mitochondrial protein synthesis blockade in curtailing growth of PDAC with fibrolytic stroma. The differential effect of fibrolytic and inert tumor stroma on PDAC growth and metabolism explain much of the controversy regarding the effects of CAFs and Col I on PDAC progression in mice^7, 10, 34–39^. Most importantly, our findings extend to human patients. We thus propose that treatments targeting DDR1-IKKβ-NF-κB-NRF2 signaling and mitochondrial biogenesis should be evaluated in prospective clinical trials that include stromal state, an important modifier of tumor growth, as an integral biomarker. A deeper understanding of whether stromal state is impacted by neoadjuvant chemotherapy and how the stromal state affects metastasis are other areas of priority for further investigation. Although our results do not apply to TNBC, they may be applicable to other desmoplastic and fibrolytic cancers.

## Methods

### Antibodies

Antibodies specific for the following antigens were used: Guinea pig anti-p62 polyclonal antibody (GP62-C, Progen), rabbit anti-NRF2 polyclonal antibody (ABclonal, A11159), rabbit anti-COL1A1 monoclonal antibody (CST, 72026), mouse anti-COL1A1 monoclonal antibody (Santa Cruz, sc-293182), rabbit anti-3/4 COL1A1 polyclonal antibody (Immunoglobe, 0217-050), mouse anti-Tim23 monoclonal antibody (Santa Cruz, sc-514463), rabbit anti-phospho-DDR1 (pTyr513) polyclonal antibody (Sigma, SAB4504671), rabbit anti-DDR1 polyclonal antibody (Abcam, ab227195), mouse anti-DDR1 monoclonal antibody (Santa Cruz, sc-390268), rabbit anti-KEAP1 monoclonal antibody (CST, 8047), rabbit anti-NF-κB p65 monoclonal antibody (CST, 8242), rabbit anti-Hitone H3 polyclonal antibody (ABclonal, A2348), rat anti-CD326 (EpCAM) monoclonal antibody (ThermoFisher, 13-5791-80), mouse anti-IKKα monoclonal antibody (Invitrogen, MA5-16157), mouse anti-actin monoclonal antibody (Sigma, A4700), rabbit anti-GFP polyclonal antibody (Molecular Probes, A-11122), chicken anti-GFP/YFP/CFP polyclonal antibody (Abcam ab13970), mouse anti-Flag monoclonal antibody (Sigma, F3165), rabbit anti-Flag polyclonal antibody (Sigma, F7425), rabbit anti-TFAM polyclonal antibody (Abcam, ab131607), rabbit anti-PGC1α polyclonal antibody (Sigma, ABE868), rabbit anti-Phospho-AMPKα (Thr172) monoclonal antibody (CST, 2535), rabbit anti-AMPKα monoclonal antibody (CST, 5832), mouse anti-6X His tag monoclonal antibody (Abcam, ab18184), rabbit anti-E-Cadherin monoclonal antibody (CST, 3195), rabbit anti-CD138/SDC1 antibody (ThermoFisher, 36-2900), mouse anti-NHE-1 monoclonal antibody (Santa Cruz, sc-136239), rabbit anti-PI3 Kinase p110γ monoclonal antibody (CST, 5405), mouse anti-ATP5A monoclonal antibody (Santa Cruz, sc-136178), mouse anti-ATP5B monoclonal antibody (Sigma, MAB3494), mouse anti-UQCRC2 monoclonal antibody (Santa Cruz, sc-390378), mouse anti-SDHB monoclonal antibody (Santa Cruz, sc-271548), rabbit anti-SDHB monoclonal antibody (CST, 92649), mouse anti-NDUFB7 monoclonal antibody (Santa Cruz, sc-365552), rabbit anti-COX1/MT-CO1 polyclonal antibody (CST, 62101), rabbit anti-αSMA polyclonal antibody (Abcam, ab5694), rabbit anti-MMP1 monoclonal antibody (Abcam, ab52631), rabbit anti-Ki67 monoclonal antibody (GeneTex, GTX16667), rabbit anti-CDC42 polyclonal antibody (ThermoFisher, PA1-092), mouse anti-HSP90 monoclonal antibody (Santa Cruz, sc-13119), rabbit anti-α-Amylase polyclonal antibody (Sigma, A8273), goat anti-cytokeratin 19 polyclonal antibody (Santa Cruz, sc-33111), rabbit anti-SOX9 polyclonal antibody (Santa Cruz, sc-20095), mouse anti-cytokeratin 18 monoclonal antibody (GeneTex, GTX105624), rabbit anti-LAIR1 polyclonal antibody (ThermoFisher, H00003903-D01P), mouse anti-Endo180/MRC2 monoclonal antibody (Santa Cruz, sc-271148), mouse anti-Integrin β1/ITGB1 antibody (Santa Cruz, sc-374429), HRP goat anti-chicken IgY antibody (Santa Cruz, sc-2428), HRP goat anti-rabbit IgG antibody (CST, 7074), HRP horse anti-mouse IgG antibody (CST, 7076), HRP streptavidin (Pharmingen, 554066), Biotin goat anti-mouse IgG (Pharmingen, 553999), Biotin goat anti-rabbit IgG (Pharmingen, 550338), Biotin mouse anti-goat IgG (Santa Cruz, sc-2489). Alexa 594-, Alexa 647-, and Alexa 488-conjugated secondary antibodies were used: donkey anti-mouse IgG, donkey anti-rabbit IgG, goat anti-chicken IgY (Molecular Probes, Invitrogen).

### Cell Culture

All cells were incubated at 37℃ in a humidified chamber with 5% CO2. MIA PaCa-2, UN-KPC-960, UN-KC-6141 cells, WT and R/R fibroblasts were maintained in DMEM (Invitrogen) supplemented with 10% fetal bovine serum (FBS) (Gibco). WT and R/R fibroblasts were generated at Dr. David Brenner lab16. 1305 primary human PDAC cells were generated by Dr. Andrew M. Lowy lab from a human PDAC PDX^17^ and were maintained in RPMI (Gibco) supplemented with 20% FBS and 1 mM sodium pyruvate (Corning). All media were supplemented with penicillin (100 mg/ml) and streptomycin (100 mg/ml).

### Plasmids

For gene ablations, the target cDNA sequences of mouse *Ddr1* (GTAACGCAACCGATAGCTTC), *Mrc2* (CCGGTGGACCAATGTCAAGG), *Itgb* (AATGTCACCAATCGCAGCAA), *Lair1* (GTCCGAACGTAGTAAGACGC), *Nrf2* (GGCATCTTGTTTGGGAATGT), *Col1α1* (CGTGCAATGCAATGAAGAAC), and human *DDR1* (GGATCTACAACGACTGCACC) were cloned into lentiCRISPR v2-Blast vector or lentiCRISPR v2-puro vector, respectively using BsmBI. For gene knockdowns (KD), pLKO.1-puro-Ddr1 (TRCN0000023369), pLKO.1-puro-DDR1 (TRCN0000121163) and pLKO.1-puro-Sdc1 (TRCN0000302270) were ordered from Sigma. pCDH-CMV-MCS-EF1-puro-Col1α1-6*His and pLVX-IRES-Puro-NRF2^E79Q^-Flag were made by Sangon Biotech (Shanghai, China). pLKO.1-blast-Ikkα, pLKO.1-puro-Nhe1, pLKO.1-puro-NHE1, pLKO.1-puro-NRF2, and lentiCRISPR v2-Puro-p62/*Sqstm1* were described^17^. LentiCRISPR v2-Blast-ATG7^40^ was a gift from Dr. Sina Ghaemmaghami.

### Stable cell line construction

To generate lentiviral particles, HEK293T cells were transfected with the above LentiCRISPR v2, pLVX-IRES-Puro or pLKO.1 vectors (7.5 μg), pSPAX2 (3.75 μg) and pMD2.G (3.75 μg) DNAs. The next day, the medium was exchanged to fresh antibiotic-free DMEM or RPMI plus 20% FBS. After 2 days, virus particle-containing media were harvested, filtered and stored in -80℃. MIA PaCa-2, 1305, KPC, KC cells or fibroblasts were transduced by combining 1 ml of viral particle-containing medium with 8 μg/ml polybrene. The cells were fed 8 h later with fresh medium and selection was initiated 48 h after transduction using 1.25 μg/ml puromycin or 10 μg/ml blasticidin. IKKα KD KC6141 cell, NRF2^KD^ MIA PaCa-2 cell and ATG7^Δ^ MIA PaCa-2 cell were described^17^.

### Mice

Female homozygous *Nu/Nu* nude mice and C57BL/6 mice were obtained at 6 weeks of age from Charles River Laboratories and The Jackson Laboratory, respectively. *Col1α1^+/+^* (Col I^WT^) or *Col1α1^r/r^* (Col I^r/r^) mice on a C57BL/6 background were obtained from Dr. David Brenner at UCSD and were previously described^13, 41^. Age- and sex-matched (except where otherwise indicated) male and female mice of each genotype were generated as littermates for use in experiments in which different genotypes were compared. For xenograft studies, female mice were randomly allocated to different treatment groups after cell injections. All mice were maintained in filter-topped cages on autoclaved food and water, and experiments were performed in accordance with UCSD Institutional Animal Care and Use Committee and NIH guidelines and regulations on age and gender-matched littermates. Dr. Karin’s Animal Protocol S00218 was approved by the UCSD Institutional Animal Care and Use Committee. The number of mice per experiment and their age are indicated in the figure legends.

### Orthotopic PDAC cell implantation

Col I^WT^ or Col I^r/r^ mice were pretreated with or without 50 μg/kg caerulein (CAE) intraperitoneal (i.p.) injections every h, 6 times daily on 1^st^, 4^th^ and 7^th^ day. On the 11^th^ day, parental, NRF2^E79Q^, DDR1^KD^, or DDR1^KD^+NRF2^E79Q^ KPC or KC cells were orthotopically injected into 3-month-old Col I^WT^ or Col I^r/r^ mice as described^17^. Briefly, mice were anesthetized with Ketamine/Xylazine (100 mg/kg and 10 mg/kg body weight, respectively). After local shaving and disinfection, a 1.5 cm long longitudinal incision was made on the left upper quadrant of abdomen. The spleen was lifted and 50 μl of 10^6^ cell suspension in ice-cold PBS-Matrigel mixture (equal amounts) was slowly injected into the tail of the pancreas. Successful injection was confirmed by formation of a liquid bleb at the site of injection with minimal fluid leakage. Following surgery, mice were given buprenorphine subcutaneously at a dose of 0.05-0.1 mg/kg every 4-6 h for 12 h and then every 6-8 h for 3 additional days. Mice were analyzed after 30 days. Throughout the experiment, animals were provided with food and water ad libitum and subjected to a 12-h dark/light cycle.

### Intrasplenic PDAC cell implantation

3-month-old Col I^WT^ or Col I^r/r^ mice were pretreated with or without an oral gavage of 25% CCl_4_ in corn oil twice a wk for 2 wk. After 2 wk recovery, parental, NHE1^KD^, or IKKα^KD^ KPC or KC cells (10^6^ cells in 50 μl PBS) were adoptively transferred into livers of Col I^WT^ or Col I^r/r^ mice via intrasplenic injection, followed by immediate splenectomy^42^. Mice were analyzed 14 days after treatment with or without 10 mg/kg EIPA (Sigma) by i.p. injection every other day. Throughout the experiment, animals were provided with food and water ad libitum and subjected to a 12-h dark/light cycle.

### Subcutaneous PDAC cell implantation

Homozygous BALB/c Nu/Nu females were injected subcutaneously (s.c.) in single or both flanks at 7 wk of age with 5x105 parental, NHE1KD, DDR1KD, or DDR1KD+NRF2E79Q MIA PaCa-2 cells or 1305 cells mixed with or without 5x10^5^ WT, R/R, Col I^Δ^ WT, Col I^Δ^ R/R fibroblasts diluted 1:1 with BD Matrigel (BD Biosciences) in a total volume of 100 μl. Tumors were collected after 4 weeks. To evaluate the effect of IKKβ or mitochondrial protein synthesis inhibition on tumor growth, mice were treated with vehicle (DMSO in PBS), ML120B (60 mg/kg) twice daily through oral gavage or tigecycline (50 mg/kg) twice daily through intraperitoneal injection for 3 weeks. Therapy was started one week after tumor implantation. Volumes (1/2 × (width^2^ × length)) of s.c. tumors were calculated based on digital caliper measurements. Mice were euthanized to avoid discomfort if tumor diameter reached 2.0 cm.

### Human specimens

Survival analysis of high and low Col-MMP patients was performed using TCGA data and the GEPIA2 platform. The collagen-cleaving signature consisted of MMP1, MMP2, MMP8, MMP9, MMP13 and MMP14. Overall survival was determined in the TCGA cohort of 178 PDAC patients using a median cut-off.

Col1α1, 3/4 Col1α1, DDR1, NF-κB p65, NRF2, NHE1, CDC42, SDC1, SDHB, α-SMA, and MMP1 protein expression in 106 human PDAC specimens was analyzed and is shown in Fig. 3a-c and Extended Data Fig. 6a, b. These tissues were acquired from patients who were diagnosed with PDAC between January 2017 and May 2021 at The Affiliated Drum Tower Hospital of Nanjing University Medical School (Nanjing, Jiangsu, China). All patients received standard surgical resection and did not receive chemotherapy before surgery. Paraffin embedded tissues were processed by a pathologist after surgical resection and confirmed as PDAC prior to further investigation. Overall survival duration was defined as the time from date of diagnosis to that of death or last known follow-up examination. Survival information was available for 81 of the 106 patients. The study was approved by the Institutional Ethics Committee of The Affiliated Drum Tower Hospital with IRB #2021-608-01. Informed consent for tissue analysis was obtained before surgery. All research was performed in compliance with government policies and the Helsinki declaration.

### Immunohistochemistry

Pancreata or liver were dissected and fixed in 4% paraformaldehyde in PBS and embedded in paraffin. 5 μm sections were prepared and stained with hematoxylin and eosin (H&E) or Sirius Red (SR). For IHC, after xylene de-paraffinization and rehydration through graded ethanol, antigen retrieval was performed for 20 min at 100°C with 0.1% sodium citrate buffer (pH 6.0). Following quenching of endogenous peroxidase activity with 3% H2O2 and blocking of non-specific binding with 5% bovine serum albumin (BSA), sections were incubated overnight at 4°C with the appropriate primary antibodies followed by incubation with 1:500 biotinylated secondary antibodies for 45 min and 1:500 streptavidin-HRP for 45 min. Bound peroxidase was visualized by 1-10 min incubation in a 3, 3’-diaminobenzidine (DAB) solution (Vector Laboratories, SK-4100). Slides were photographed on an upright light/fluorescent Imager A2 microscope with AxioVision Release 4.5 software (Zeiss, Germany).

### IHC Scoring

A modified labeling score (H score) which is calculated by using percentage of positive stained cancer cells and their intensity per tissue core^17^ was used. Staining intensity was determined by dominant staining and was divided into four categories (0, negative; 1, weak; 2, moderate; 3, strong). By multiplying staining intensity and percentage of positive stained tumor cells ranging from 0 to 100%, H score recieved a range of 0-300. Cores with overall scores of 0-5, 5-100, 101-200, 201-300 were classified as negative, weak, intermediate and strong expression level. Complete absence of staining or less than 5% of cancer cells stained faintly were considered to be negative (H-score, 0-5). Negative and weak were viewed as low expression level and intermediate and strong were viewed as high expression level. For cases with tumors with two satisfactory cores, the results were averaged; for cases with tumors with one poor-quality core, results were based on the interpretable core. Based on this evaluation system, Chi-square test was used to estimate the association between Col I-DDR1-NRF2 signaling proteins staining intensities. The number of evaluated cases for each different staining in PDAC tissues and the scoring summary are indicated (Extended Data Fig. 6a).

### Extracellular matrix (ECM) preparation

WT or R/R fibroblasts were seeded on 6/12/96-well plates. One day after plating, cells were switched into DMEM (with pyruvate) with 10% dialyzed FBS supplemented with or without 500 μM ^3^H-proline or [U-^13^C]glutamine and 100 μM vitamin C. Cells were cultured for 5 days with media renewal every 24 h. Then fibroblasts were removed by washing in 1 ml or 500 μl or 100 μl per well PBS with 0.5% (v/v) Triton X-100 and 20 mM NH4OH. The ECM was washed five times with PBS before cancer cell plating. The following day, cancer cells were switched into the indicated medium for 24 or 72 h.

### Cell imaging

Immunostaining was performed as described^43^. Cells were cultured on coverslips coated with or without ECM and fixed in 4% PFA for 10 min at room temperature or methanol for 10 min at -20℃. After washing twice in PBS, cells were incubated in PBS containing 10% FBS to block nonspecific sites of antibody adsorption. The cells were then incubated with the appropriate primary antibodies (diluted 1:100) and secondary antibodies (diluted 1:500) in PBS containing 10% FBS with or without 0.1% saponin. Macropinosomes were imaged using a high-molecular-mass (70 KDa) TMR-DEX (Invitrogen)-uptake assay, wherein TMR-DEX was added to serum-free medium at a final concentration of 1 mg/ml for 30 min at 37℃. At the end of the incubation period, the cells were rinsed five times in cold PBS and immediately fixed in 4% PFA. For tissue macropinosome visualization^44^, fresh pancreata were cut into pieces of an approximate 5-mm cuboidal shape. Tissue fragments were placed in a 24-well plate and injected with 150 μL of 10 mg/ml TMR-DEX solution. Another 250 μL of diluted TMR-DEX solution were added to the well to immerse the tissue fragment. The plate was incubated in the dark for 15 min at room temperature. Then the tissue fragments were rinsed twice in PBS before embedding in O.C.T. compound in a prelabeled cryomold. Specimens were frozen on dry ice and stored at -80℃ for further processing. Immunostaining was performed as described above. Images were captured using a TCS SPE Leica confocal microscope (Leica, Germany).

### Second harmonic generation (SHG)

Mouse pancreatic tumor tissue was fixed in 4% paraformaldehyde in PBS and embedded in paraffin. 5 μm sections were prepared and deparaffinized in xylene, rehydrated in graded ethanol series as described^45^, mounted using an aqueous mounting medium and sealed with a coverslip. All samples were imaged using a Leica TCS SP5 multiphoton confocal microscope and a HC APO LC 20× 1.00W was used throughout the experiment. The excitation wavelength was tuned to 840 nm, and a 420±5 nm narrow bandpass emission controlled by a prism was used for detecting the SHG signal of collagen. SHG signal is generated when two photons of incident light interact with the non-centrosymmetric structure of collagen fibers, which leads to the resulting photons being half the wavelength of the incident photons. SHG measurements were performed using CT-Fire software (v.2.0 beta) (https://loci.wisc.edu/software/ctfire). Tumor area was confirmed by H&E staining.

### Immunoblot and immunoprecipitation

Cells were harvested and lysed in RIPA buffer (50 mM Tris-HCl, pH 7.4, 150 mM NaCl, 1% Triton X-100, 1% Na deoxycholate, 0.1% SDS, 1 mM EDTA) supplemented with complete protease inhibitor cocktail. Proteins were resolved on SDS-polyacrylamide gels, and then transferred to a polyvinylidene difluoride membrane. After blocking with 5% (w/v) fat free milk or 5% BSA, the membrane was stained with the corresponding primary antibodies followed by incubation with the appropriate secondary HRP-conjugated antibodies, and development with ECL. Immunoreactive bands were detected by automatic X-ray film processor or a KwikQuant Imager.

For immunoprecipitation, cells were lysed in Nonidet P-40 (NP-40) lysis buffer (20 mM Tris-HCl, pH 7.5, 1% NP-40, 137 mM NaCl, 1 mM MgCl2, 1 mM CaCl2, 10% glycerol) supplemented with a complete protease inhibitor cocktail. Immunoprecipitation was performed using indicated antibodies. Generally, 2 mg of antibody were added to 1 mL of cell lysate and incubated at 4℃ overnight. After addition of protein A/G-magnetic beads, incubation was continued for 2 h, and then immune complexes were washed five times using lysis buffer, resolved by SDS-PAGE, and analyzed by immunoblot.

### Chromatin immunoprecipitation (ChIP)

Cells were crosslinked with 1% formaldehyde for 10 min and the reaction was stopped with 0.125 M glycine for 5 min. ChIP assay was performed as described^17^. Cells were lysed and sonicated on ice to generate DNA fragments with an average length of 200-800 bp. After pre-clearing, 1% of each sample was saved as the input fraction. IP was performed using antibodies specifically recognizing NRF2 (CST, 12721). DNA was eluted and purified from complexes, followed by PCR amplification of the target promoters or genomic loci using primers for mouse *Tfam*: 5′-GAGGCAGGGTCTCATG-3′, and 5′-CAAGCTGAGTTCTATC-3′; 5′-TCTGGGCCATCTTGGG-3′, and 5′-CCATGGGCCTGGGCTG-3′.

### Real-time PCR analysis

Total RNA and DNA were extracted using All Prep DNA/RNA Mini Kit (Qiagen). RNA was reverse transcribed using a Superscript VILO cDNA synthesis kit (Invitrogen). Real-time PCR (RT-PCR) was performed as described^15^. Relative expression levels of target genes were normalized against the level of 18s rRNA expression. Fold difference (as relative mRNA expression) was calculated by the comparative CT method (2^Ct(18s rRNA-gene of interest)^). Primers obtained from the NIH Primer-BLAST (https://www.ncbi.nlm.nih.gov/tools/primer-blast/index.cgi?LINK_LOC=BlastHome) were as follows: *Pik3ca* F: 5′-GGACTGTGTGGGTCTCATCG-3′; *Pik3ca* R: 5′-TCTCGCCCTTGTTCTTGTCC-3′; *Pik3cg* F: 5′-CTCTG GACCTGTGCCTTCTG-3′; *Pik3cg* R: 5′-ATCTTTGAATGCCCCCGTGT-3′; *Cdc42* F: 5′-GAGACTGCTGAAAAGCTGGCG-3′; *Cdc42* R: 5′-GGCTCTTCTTCGGTTCTGGAGG-3′; *Nfe2l2* F: 5’-AACAGAACGGCCCTAAAGCA-3’; *Nfe2l2* R: 5′-GGGATTCACGCATAGG AGCA-3’; *Nhe1* F: 5′-TCATGAAGATAGGTTTCCATGTGAT-3′; *Nhe1* R: 5′-CGTCTGA TTGCAGGAAGGGG-3′; *Sdc1* F: 5′-TCTGGCTCTGGCTCTGCG-3′; *Sdc1* R: 5′-GCCGTGA CAAAGTATCTGGC-3′; *Sqstm1* F: 5’-TGGGCAAGGAGGAGGCGACC-3’; *Sqstm1* R: 5’-CCTCATCGCGGTAGTGCGCC-3’; *Egf* F: 5′-TTCTCACAAGGAAAGAGCATCTC-3′; *Egf* R: 5′-GTCCTGTCCCGTTAAGGAAAAC-3′; *m18s* F: 5′-AGCCCCTGCCCTTTG TACACA-3′; *m18s* R: 5′-CGATCCGAGGGCCTCACTA-3′.

### RNA-seq library preparation, processing and analysis

Total RNA was isolated as described above from KPC samples grown on WT (n=3) or R/R (n=3) ECM as indicated. RNA purity was assessed by an Agilent 2100 Bioanalyzer. 500 ng total RNA was enriched for poly-A tailed RNA transcripts by double incubation with Oligo d(T) Magnetic Beads (NEB, S1419S) and fragmented for 9 min at 94°C in 2X Superscript III first-strand buffer containing 10 mM DTT (Invitrogen, #P2325). Reverse-transcription (RT) reaction was performed at 25°C for 10 min followed by 50°C for 50 min. RT product was purified with RNAClean XP (Beckman Coulter, #A63987). Libraries were ligated with dual UDI (IDT) or single (Bioo Scientific), PCR-amplified for 11-13 cycles, size-selected using one-sided 0.8× AMPure cleanup beads, quantified using the Qubit dsDNA HS Assay Kit (Thermo Fisher Scientific), and sequenced on a HiSeq 4000 or NextSeq 500 (Illumina).

RNA-seq reads were aligned to the mouse genome (GRCm38/mm10) using STAR. Biological and technical replicates were used in all experiments. Quantification of transcripts was performed using HOMER. Principal Component Analysis (PCA) was obtained based on Transcripts Per kilobase Million (TPM) on all genes from all samples. Expression value for each transcript was calculated using the analyzeRepeats.pl tool of HOMER. Differential expression analysis was calculated using getDiffExpression.pl tool of HOMER. Pathway analyses were performed using the Molecular Signature Database of GSEA.

### Single cell RNA-seq

PDAC patient samples from five primary tumors and one PDAC liver metastasis were obtained from a published dataset^46^ and their method was followed to filter low quality cells and obtain batch-corrected integrated samples. Briefly, the dataset was processed in R (v4.0.2) and Seurat^47^ (v4.0.5) and cells with at least 200 genes and genes expressed in at least 3 cells were retained for further analysis. The resulting gene-cell barcode matrix was log-normalized, 2000 variable genes identified and scaled to perform principal component analysis (PCA). Individual samples were batch-corrected and integrated using reciprocal PCA (RPCA) pipeline in Seurat to obtain a combined sample of nine PDAC primary and metastatic samples. After obtaining this batch-corrected integrated sample, we obtained the primary PDAC samples (P1, P2, P3, P4, P5) using ‘subset’ function. The resulting integrated expression matrix was scaled and PCA was performed. To cluster and visualize the cells, ‘FindNeighbours’, ‘FindClusters’ and ‘RunUMAP’ functions were used on top 12 principal components which were selected using ‘ElbowPlot’. The PDAC liver-metastasis sample was also obtained from the above batch-corrected integrated sample using ‘subset’ function and followed identical steps of primary PDAC samples to obtain cell clusters and visualization.

The cell types were identified by manual annotation of the well-known makers^46^, namely: epithelial-tumor cells (*EPCAM*, *KRT8*), T cell (*CD3D*, *IL7R*), Myeloid (*CD14*, *LYZ*), NK (*FCGR3A*, *NKG7*, *GNLY*), B cell (*CD79A*, *MS4A1*), dendritic cell (*TSPAN13*, *GPR183*), endothelial cells (*PECAM1*, *CDH5*), fibroblasts (*ACTA2*, *COL1A1*), hepatocytes (*ALB*, *APOB*, *CPS1*), cholangiocytes (*ANXA4*, *KRT7*, *SOX9*), plasma cells (*JCHAIN*, *IGKC*) and cycling cells (*TOP2A*, *MKI67*).

### Metabolite extraction and analysis

Cells grown on a 12-well plate coated with or without ECM were rinsed with 1 ml cold saline and quenched with 250 μl cold methanol. 100 μl of cold water containing 1 μg norvaline was added, cell lysates were collected, and 250 μl of chloroform was added to each sample. After extraction the aqueous phase was collected and evaporated under nitrogen.

Dried polar metabolites were dissolved in 2% methoxyamine hydrochloride in pyridine (Thermo) and held at 37℃ for 1.5 h. After dissolution and reaction, tertbutyldimethylsilyl derivatization was initiated by adding 30 ml N-methyl-N-(tertbutyldimethylsilyl) trifluoroacetamide + 1% tert-butyldimethylchlorosilane (Regis) and incubating at 37℃ for 1 h. GC/MS analysis was performed using an Agilent 6890 GC equipped with a 30m DB-35MS capillary column connected to an Agilent 5975B MS operating under electron impact ionization at 70 eV. 1 μl of sample was injected in splitless mode at 270℃, using helium as the carrier gas at a flow rate of 1 ml/min. For measurement of amino acids, the GC oven temperature was held at 100℃ for 3 min and increased to 300℃ at 3.5℃/min. The MS source and quadrupole were held at 23℃ and 150℃, respectively, and the detector was run in scanning mode, recording ion abundance in the range of 100-605 *m/z*. Mole percent enrichments of stable isotopes in metabolite pools were determined by integrating the appropriate ion fragments and correcting for natural isotope abundance as previously described^48^.

### Cell viability assay

Cell viability was determined with Cell Counting Kit-8 assay (CCK-8 assay kit, Glpbio). Cells were plated in 96-well plates coated with or without ECM at a density of 3000 cells (MIA PaCa-2, 1305) or 1500 cells (KPC or KC6141) per well and incubated overnight prior to treatment. 7rh (500 nM), ML120B (10 μM), EIPA (10.5 μM), IPI549 (600 nM), MBQ-167 (500 nM), MRT68921 (600 nM) or, ML385 (10 μM, Sigma), or their combinations were added to the wells in the presence of complete medium (CM), low glucose medium (LG; Glucose-free DMEM medium supplemented with 0.5 mM glucose in the presence of 10% dialyzed FBS and 25 mM HEPES) or low glutamine medium (LQ; Glutamine-free DMEM medium was supplemented with 0.2 mM glutamine in the presence of 10% dialyzed FBS and 25 mM HEPES) for 72 h. Next, 10 μL of CCK-8 was added to each well. Optical density was read at 450 nm using a microplate reader (FilterMax F5, Molecular Devices, USA). For all experiments, media were replaced every 24 h.

### Luminescence ATP detection assay

Intracellular ATP was determined with luminescence ATP detection assay system (PerkinElmer) according to manufacturer’s protocol. Briefly, KPC or KC cells were grown on 96-well plates coated with or without indicated ECM in the presence of 100 μl CM or LG medium with or without EIPA (10.5 μM), MBQ-167 (500 nM), MRT68921 (600 nM) or their combinations for 24 h. 50 μl mammalian cell lysis solution was added and shaken for 5 min. Next, 50 μl substrate solution was added and shaken for 5 min. Finally, the plate was adapted in dark for 10 min and luminescence was measured.

### L-amino acid assay

Total amounts of free L-amino acids (except for glycine) were measured by L-Amino Acid Assay Kit (Colorimetric) (antibodies) according to manufacturer’s protocol. Briefly, KPC or KC cells were grown on 6-well plates coated with or without indicated ECM in the presence of 100 μl LG medium with or without EIPA (10.5 μM), MRT68921 (600 nM) or their combinations for 24 h. The cells were resuspended at 10^6^ cells/mL 1xAssay Buffer and were homogenized on ice and then centrifuged to remove debris. 50 μL of each L-Alanine standard or cell lysates were added into wells of a 96-well microtiter plate. Then 50 μL of Reaction Mix were added to each well. The well contents were mixed thoroughly and incubated for 90 min at 37°C protected from light. The plate was read with a spectrophotometric microplate reader in the 540-570 nm range. The concentration of L-Amino Acids was calculated within samples by comparing the sample OD to the standard curve.

### Quantification and statistics

Macropinosomes or mitochondria were quantified by using the ‘Analyze Particles’ feature in Image J (National Institutes of Health). Macropinocytotic uptake index^49^ or mitochondria number was computed by macropinosome or mitochondria area in relation to total cell area for each field and then by determining the average across all the fields (6 fields). These measurements were done on randomly-selected fields of view. Two-tailed unpaired Student’s t test was performed for statistical analysis using GraphPad Prism software. Data are presented as mean ± SEM. Kaplan-Meier survival curves were analyzed by log rank test. Statistical correlation between Col I-DDR1-NRF2 signaling proteins in human PDAC specimens was determined by Chi-square test. (****p < 0.0001, ***p < 0.001, **p < 0.01, *p < 0.05).

## Acknowledgments

We thank M.K. lab members for helpful discussions, Cell Signaling Technologies, Santa Cruz Technologies, ThermoFisher, Promega, and MedChemExpress for gifts of antibodies and reagents, UCSD Tissue Technology Shared Resources (TTSR) and Jennifer Santini at the UCSD School of Medicine Microscopy Core for histology and microscopy services. Research was supported by grants from the Padres Pedal the Cause/C3 (PPTC2018 to M.K., and A.M.L.), the Youth Program of the National Natural Science Foundation of China (81802757 to H.S. and 82002931 to F.Y.), the NIH (R01CA211794, R37AI043477, P01DK098108 to M.K., R01CA155630 to A.M.L., R01CA234245, R01CA218254 to C.M.M.), the National Key Research and Development Program of China (2016YFC0905900 to B.S.), the CCSG (P30CA23100 to TTSR), and UCSD School of Medicine Microscopy Core (NINDS P30-NS047101). Additional support was provided by Ride the Point (M.K. and A.M.L.), Research for a Cure of Pancreatic Cancer Fund and the Alexandrina M. McAfee Trust Foundation (A.M.L.), the UC Pancreatic Cancer Consortium to A.M.L. who is the Homer T. Hirst III Professor of Oncology in Pathology and M.K. who is the Ben and Wanda Hillyard Chair for Mitochondrial and Metabolic Diseases.

## Author contributions

M.K. and H.S. conceived the project. H.S. designed the study and H.S. and F. Y. performed most experiments. F.Y., H.S., and R.F. performed IHC analysis of human and mouse PDAC. H.S., F.Y., B.T., and J.S. performed qPCR analysis. C.M.M. and A.K. carried out the ^13^C-tracing experiments. A.M., N.S. and S.D. performed Seahorse experiments. J.L. performed several orthotopic PDAC cell implantation. J.B. performed several i.s. PDAC implantations. R.F.S., and A.N. performed single cell RNAseq and analysis. S.B.R. performed RNAseq analysis. D.B. provided Col I^r/r^ mice. B.S. collected human PDAC tissue, supervised F.Y. and R.F., and supported F.Y., and R.F. M.K. and H.S. wrote the manuscript, with all authors contributing and providing feedback and advice.

---

**Extended Data Fig. 1:**
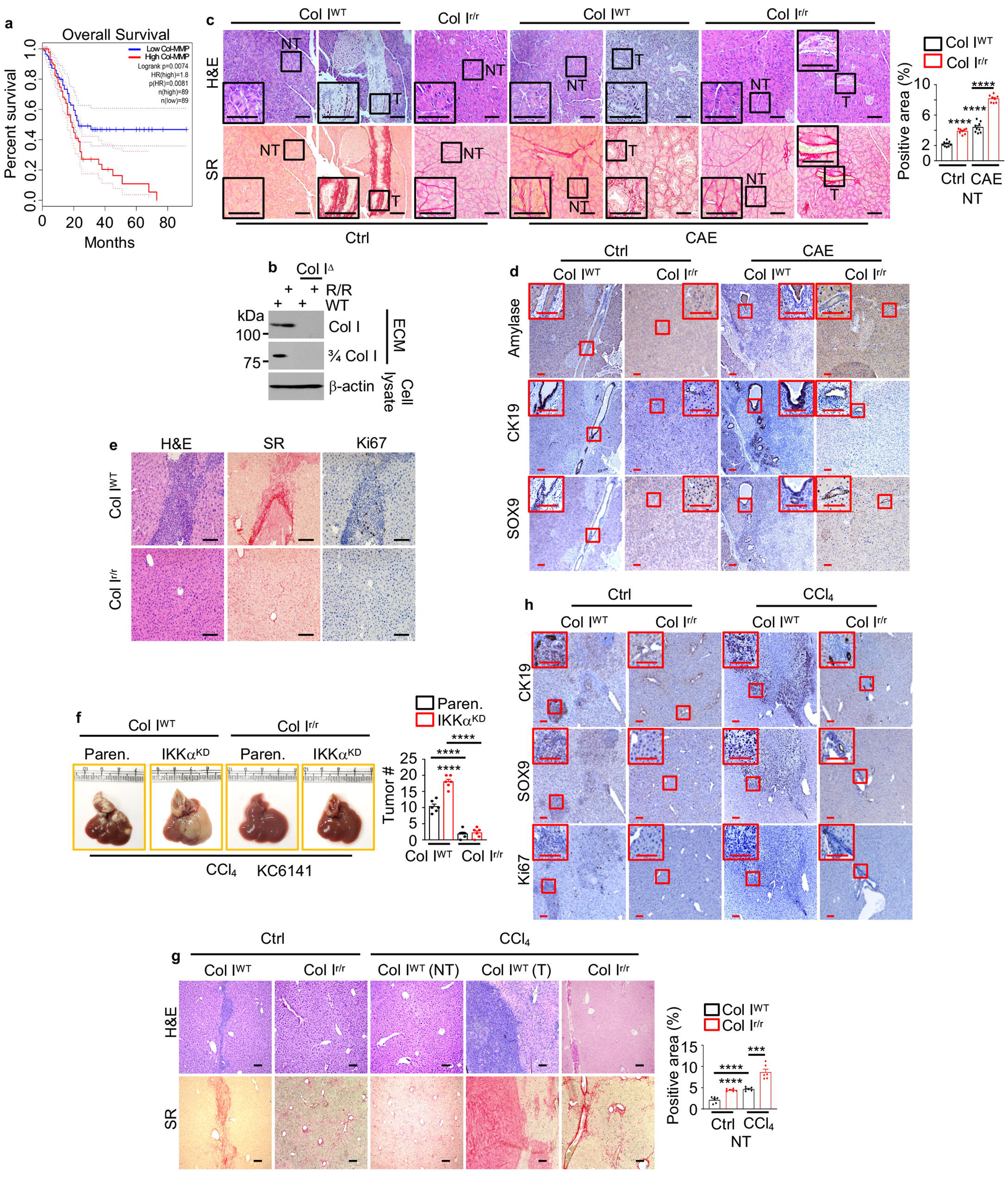
The Col I state positively and negatively controls tumor growth. **a,** Overall survival of PDAC patients from TCGA with high and low collagen-cleaving MMP signature (MMP1, 2, 8, 9, 13, 14). Significance was analyzed by log rank test. **p < 0.01. **b,** immunoblot (IB) analysis showing specificity of antibodies to iCol I and cCol I (3/4 Col I) in the ECM produced by the indicated fibroblasts. **c**, H&E and Sirius Red (SR) staining of Col I^WT^ and Col I^r/r^ pancreata 4 wk after orthotopic transplantation of KPC cells into mice pretreated -/+CAE. The boxed areas were further magnified. Quantification of SR positivity in nontumor (NT) areas is shown to the right. **d**, Immunohistochemical (IHC) analysis of pancreatic sections from above mice. **e**, H&E, SR, Ki67 staining of liver sections from above CAE-pretreated mice. **f**, Liver gross morphology and tumor numbers (#) 2 wk after i.s. transplantation of parental (Paren.) or IKKα knock down (KD) KC cells into CCl_4_ pretreated Col I^WT^ or Col I^r/r^ mice. **g**, H&E and SR staining of liver sections 2 wk after i.s. transplantation of KPC cells into Col I^WT^ and Col I^r/r^ mice pretreated -/+ CCl_4_. Quantification of SR positivity in NT areas is shown to the right. **h**, IHC analysis of liver sections from above mice. Boxed areas show higher magnification. Results in (**c**) (n=8-9), (**f**) and (**g**) (n=6) are mean ± SEM. Statistical significance was determined by a two-tailed t test. ***p < 0.001, ****p < 0.0001. (**c**-**e**, and **g**, **h**) Scale bars, 100 μm.

**Extended Data Fig. 2:**
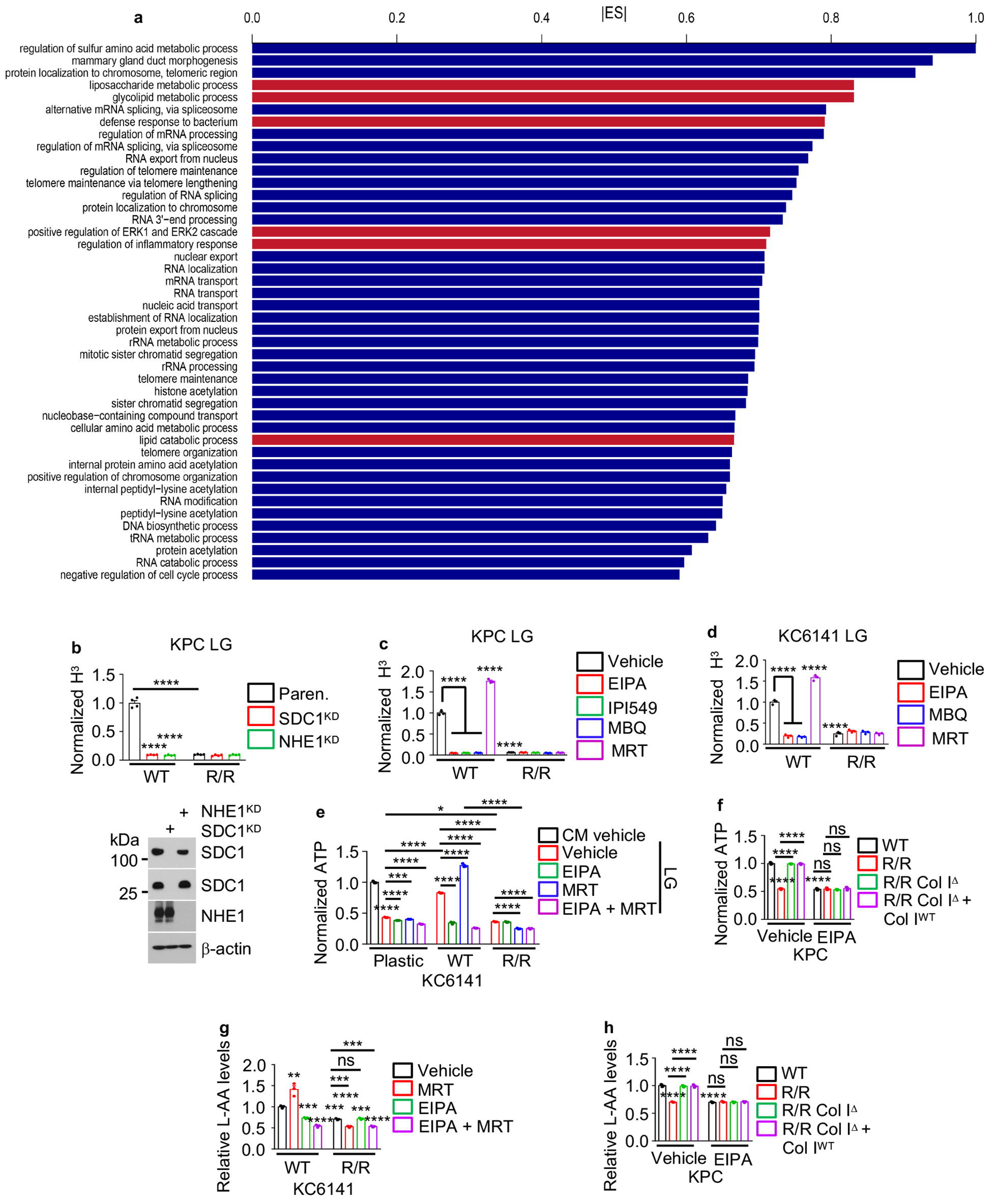
The Col I state positively and negatively controls PDAC gene expression and metabolism. **a**, Dataset enrichment graphs of RNAseq data (n=3) from KPC cells plated on WT (blue) or R/R (red) ECM and incubated in low glucose (LG, 0.5 mM) for 24 h. **b**, Paren., SDC1^KD^, or NHE1^KD^ KPC cells were plated on ^3^H-proline-labeled WT or R/R ECM and incubated in LG for 24 h. ^3^H uptake was measured by liquid scintillation counting, data were normalized to cell number and are presented as ^3^H CPM relative to Paren. KPC cells grown on WT ECM. An IB demonstrating knockdown efficiency is shown on the bottom. **c**, KPC cells were plated on ^3^H-proline-labeled WT or R/R ECM and incubated in LG -/+ EIPA (10.5 μM), IPI549 (600 nM), MBQ-167 (MBQ, 500 nM), or MRT68921 (MRT, 600 nM) for 24 h. ^3^H uptake was measured, normalized, and presented as above. **d**, ^3^H uptake by KC cells treated as above. **e**, KC cells were plated on plastic, WT or R/R ECM and incubated in complete medium (CM) or LG -/+ EIPA, MRT, or EIPA+MRT for 24 h. Total cellular ATP was measured, normalized to cell number, and presented relative to untreated plastic plated WT cells. **f**, KPC cells were plated on WT, R/R, Col I^Δ^ R/R or Col I^Δ^ RR + Col I^WT^ ECM and incubated in LG -/+ EIPA for 24 h. Total cellular ATP was measured, normalized, and presented as above. **g**, Total AA in KC cells plated on WT or R/R ECM and incubated in LG -/+ EIPA, MRT or EIPA + MRT for 24 h. Data were normalized to cell number and presented relative to untreated WT ECM plated cells. **h**, Total AA in KPC cells treated as in (**f**). Data were normalized and presented as above. Results in (**b**) (n=4 independent experiments), (**c-h**) (n=3 independent experiments) are mean ± SEM. Statistical significance was determined by a two-tailed t test. *p < 0.05, **p < 0.01, ***p < 0.001, ****p < 0.0001.

**Extended Data Fig. 3:**
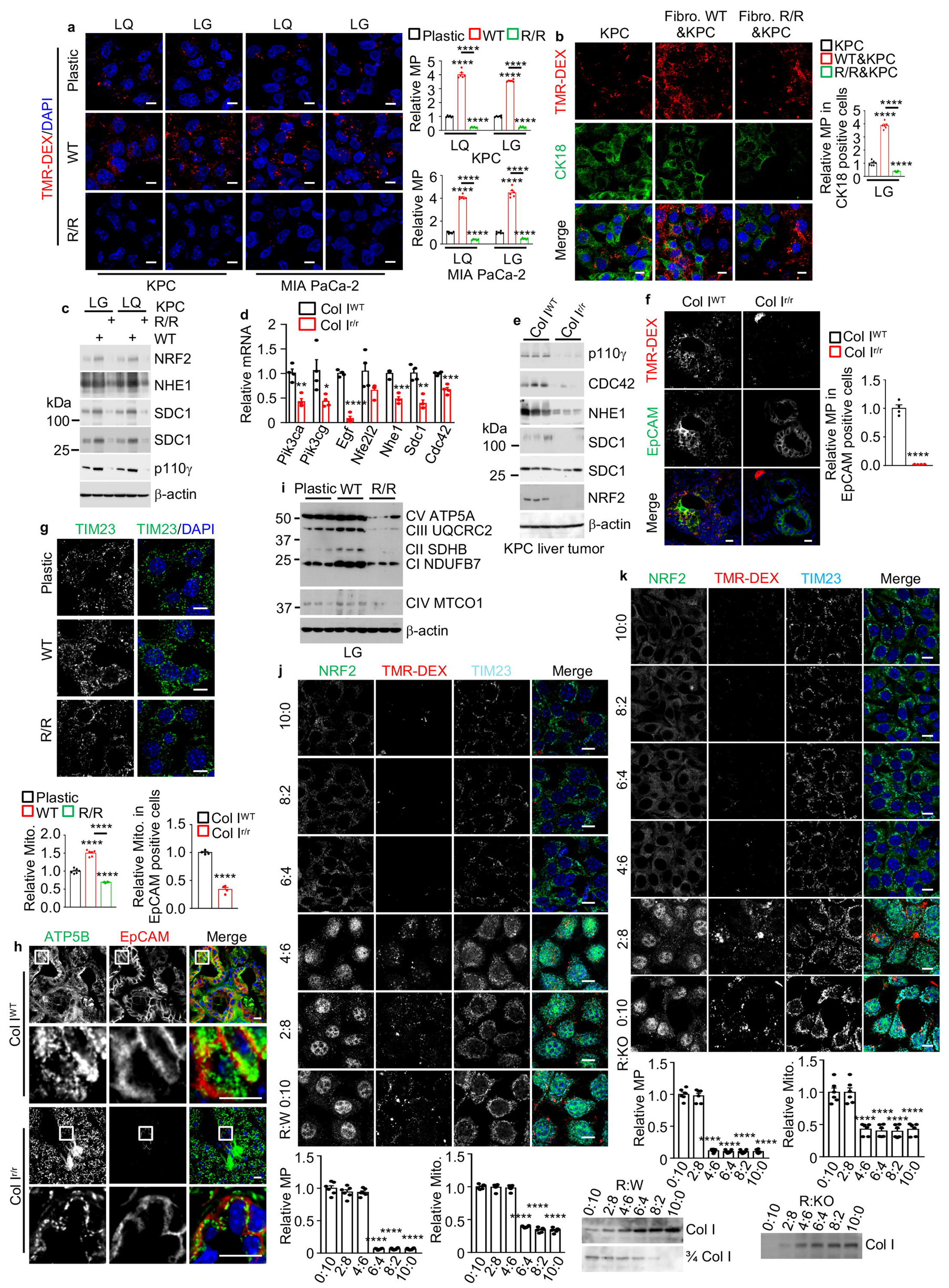
Col I state controls MP activity and mitochondria number in PDAC. **a**, Representative images, and quantification of MP in TMR-dextran (TMR-DEX, red)-incubated KPC and MIA PaCa-2 cells grown on plates +/- WT or R/R ECM and incubated in low glutamine (LQ, 0.2 mM) or LG media for 24 h. **b**, MP visualization and quantification using TMR-DEX in KPC cells cocultured with WT or R/R fibroblasts and incubated in LG medium for 24 h. KPC cell was marked by cytokeratin 18 (CK18, green). **c**, IB analysis of indicated proteins in KPC cells grown on +/- WT or R/R ECM and incubated in LG or LQ media for 24 h. **d**, qRT-PCR analysis of MP-related mRNAs of tumors isolated from liver 2 wk after i.s. transplantation of KPC cells into Col^WT^ or Col^r/r^ mice pretreated with CCl_4_. **e**, IB analysis of MP-related proteins in liver tumors isolated from above mice. **f**, Representative images and quantification of MP in TMR-DEX-injected pancreatic tissue from Col I^WT^ or Col I^r/r^ mice 4 wk after orthotopic transplantation of KPC cells. Carcinoma cells are marked by EpCAM staining (green). Quantification is on the right. Scale bars, 10 μm. Mean ± SEM (n = 4 mice). **g**, Representative images and quantification of mitochondria (Mito., TIM23) in KPC cells grown on plates +/- WT or R/R ECM and incubated in LG medium for 24 h. **h**, Representative images and quantification of Mito. (ATP5B, green) in pancreatic tissue from the indicated mice analyzed as in (**f**). Carcinoma cells are marked by EpCAM staining (red). Quantification is shown above. **i**, IB analysis of indicated proteins in KPC cells treated as in (**g**). **j**, **k**, Representative images and quantification of Mito. and MP in TMR-DEX-incubated KPC cells grown on mixed ECM produced by R/R and WT fibroblasts (R:W) (**j**) or R/R and Col I^Δ^ R/R fibroblasts (R:KO) (**k**) in the indicated ratios and incubated in LG medium for 24 h. IB analysis of iCol I or cCol I (3/4 Col I) in the above ECM preparations is shown below. Results in (**a**, **b**, **g**, **j**, **k**) (n=6), (**d**, **f**, **h**) (n=4) are mean ± SEM. Statistical significance was determined by a two-tailed t test. *p < 0.05, **p < 0.01, ***p < 0.001, ****p < 0.0001. Scale bars, 10 μm.

**Extended Data Fig. 4:**
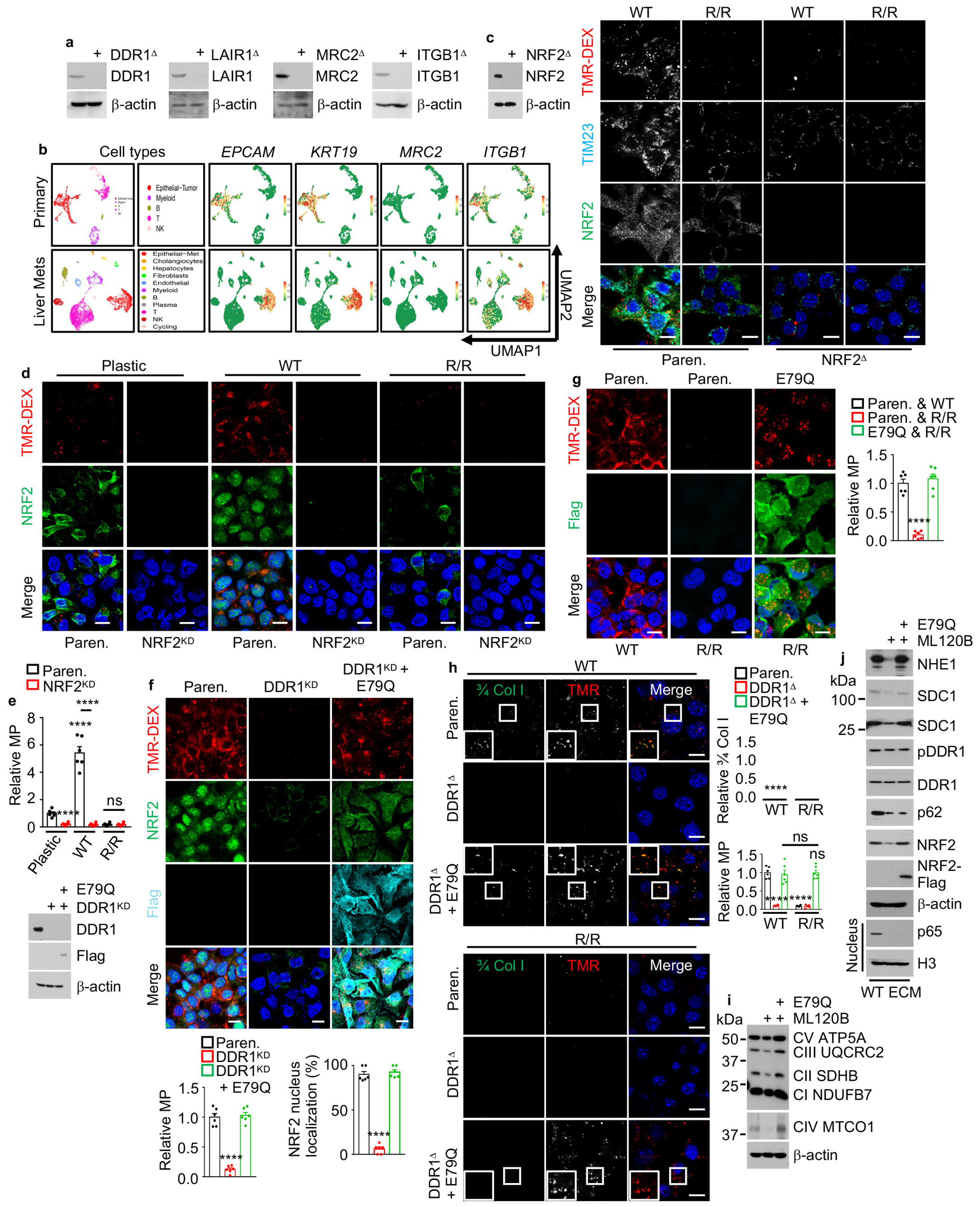
Col I controls MP and mitochondrial content through DDR1-NRF2 signaling. **a**, IB analysis of KPC cells ablated for the indicated collagen receptors. **b**, UMAPs showing scRNA-seq data from 5 patients with primary PDAC (upper row) and 1 patient with PDAC liver metastasis (lower row), displaying the identified cell populations and expression of *EPCAM*, *KRT19*, *MRC2* and *ITGB1* mRNAs. **c**, Representative images of Mito. (TIM23) and MP in TMR-DEX-incubated Paren. and NRF2^Δ^ KPC cells plated on WT or R/R ECM and incubated in LG for 24 h. IB analysis of NRF2 expression is shown on the left. **d**, MP and NRF2 localization in Paren. and NRF2^KD^ MIA PaCa-2 cells plated on plastic, WT or R/R ECM and incubated in LG for 24 h. **e**, Quantification of MP in cells from (**d**). **f**, Representative images, and quantification of MP and nuclear NRF2 in indicated 1305 cells plated on WT ECM and incubated in LG for 24 h. IB analysis of DDR1 and Flag-tagged NRF2^E79Q^ is shown on the left. **g**, MP imaging and quantification in Paren. and NRF2^E79Q^ MIA PaCa-2 cells plated on WT or R/R ECM and incubated in LG for 24 h. **h**, 3/4 Col I and MP imaging and quantification in indicated KPC cells treated as above. Although NRF2^E79Q^ stimulates MP, cCol I uptake is detected only in cells plated on WT ECM. **i**, **j**, IB analysis of indicated proteins in Paren. or NRF2^E79Q^ KPC cells plated on WT ECM and incubated in LG medium -/+ML120B (10 μM) for 24 h. ETC complex I-V (CI-CV). Results in (**e**-**h**) (n=6) are mean ± SEM. Statistical significance was determined by a two-tailed t test. ****p < 0.0001. (**c**, **d**, **f**-**h**) Scale bars, 10 μm.

**Extended Data Fig. 5:**
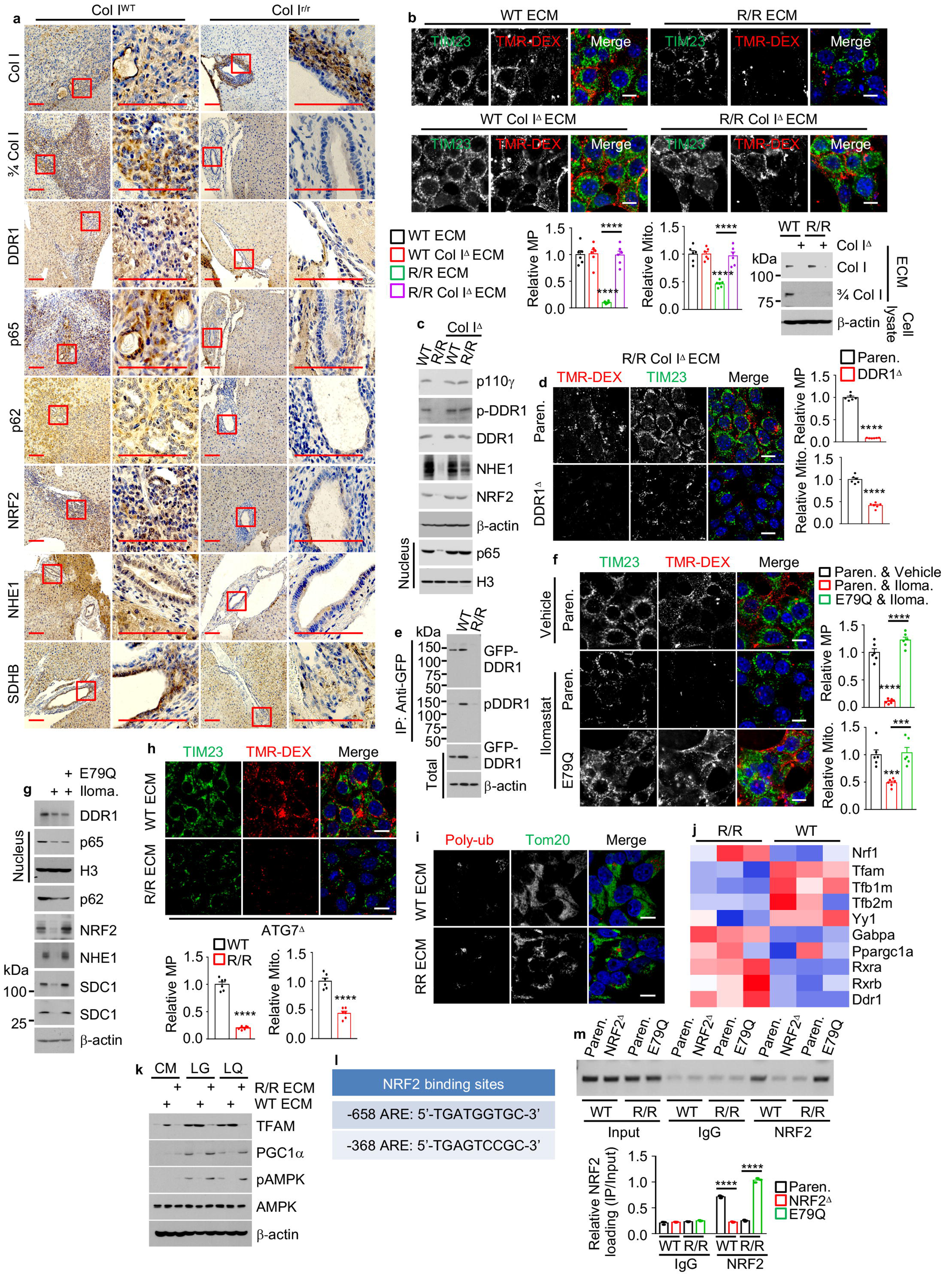
MMP inhibition exerts the same effect as intact Col I and NRF2 transcriptionally controls mitochondrial biogenesis. **a**, IHC analysis of liver sections stained as indicated and prepared 2 wk after i.s. transplantation of KPC cells into CCl_4_ pretreated Col I^WT^ and Col I^r/r^ mice. Scale bars, 100 μm. **b**, Imaging and quantification of MP and Mito. in KPC cells plated on WT, R/R, WT Col I^Δ^, or R/R Col I^Δ^ ECM and incubated in LG for 24 h. IB analysis of intact and 3/4 Col I in the ECM produced by WT or R/R fibroblasts +/- Col I^Δ^ is shown. **c**, IB analysis of indicated proteins in above cells. **d**, Imaging and quantification of MP and Mito. in Paren. and DDR1^Δ^ KPC cells plated on R/R Col I^Δ^ ECM and incubated in LG for 24 h. **e**, Immunoprecipitation (IP) analysis of GFP-DDR1 expressed in 1305 cells plated on plastic, WT or R/R ECM. **f**, Imaging and quantification of MP and Mito. in Paren. or E79Q KPC cells plated on ECM produced by Ilomastat (Iloma.) treated or untreated WT fibroblasts and incubated in LG medium -/+ Iloma. for 24 h. **g**, IB analysis of indicated proteins in above cells. **h**, Imaging and quantification of MP and Mito. in ATG7^Δ^ MIA PaCa-2 cells plated on WT or R/R ECM and incubated in LG for 24 h. **i**, Imaging of 1305 cells plated on WT or R/R ECM showing rare colocalization of poly-Ub (red) and Mito. (Tom20, green). **j**, Differentially expressed genes between KPC cells plated on WT or R/R ECM and incubated in LG for 24 h. Vertical rows: different specimens; horizontal rows: individual genes, colored according to log-transformed transcript intensity in z-scored units. TPM (transcripts per million). Blue: replicates with low expression (z-score=−2); red: replicates with high expression (z-score=+2). **k**, IB analysis of KPC cells plated on plastic, WT or R/R ECM and incubated in complete (CM), LG or LQ media for 24 h. **l**, Locations of putative NRF2 binding sites (AREs) relative to the transcriptional start site (TSS, +1) of the mouse *Tfam* gene. **m**, Chromatin IP assays probing NRF2 recruitment to the *Tfam* promoter in Paren., NRF2^Δ^, and NRF2^E79Q^ KPC cells plated on WT or R/R ECM and incubated in LG for 24 h. The image shows PCR-amplified promoter DNA fragments containing NRF2 binding sites. Quantitation is on the bottom. Results in (**b**, **d**, **f**, **h**) (n=6 fields), (**m**) (n=3 independent experiments) are mean ± SEM. Statistical significance was determined by a two-tailed t test. ***p < 0.001, ****p < 0.0001. Scale bars, 10 μm.

**Extended Data Fig. 6:**
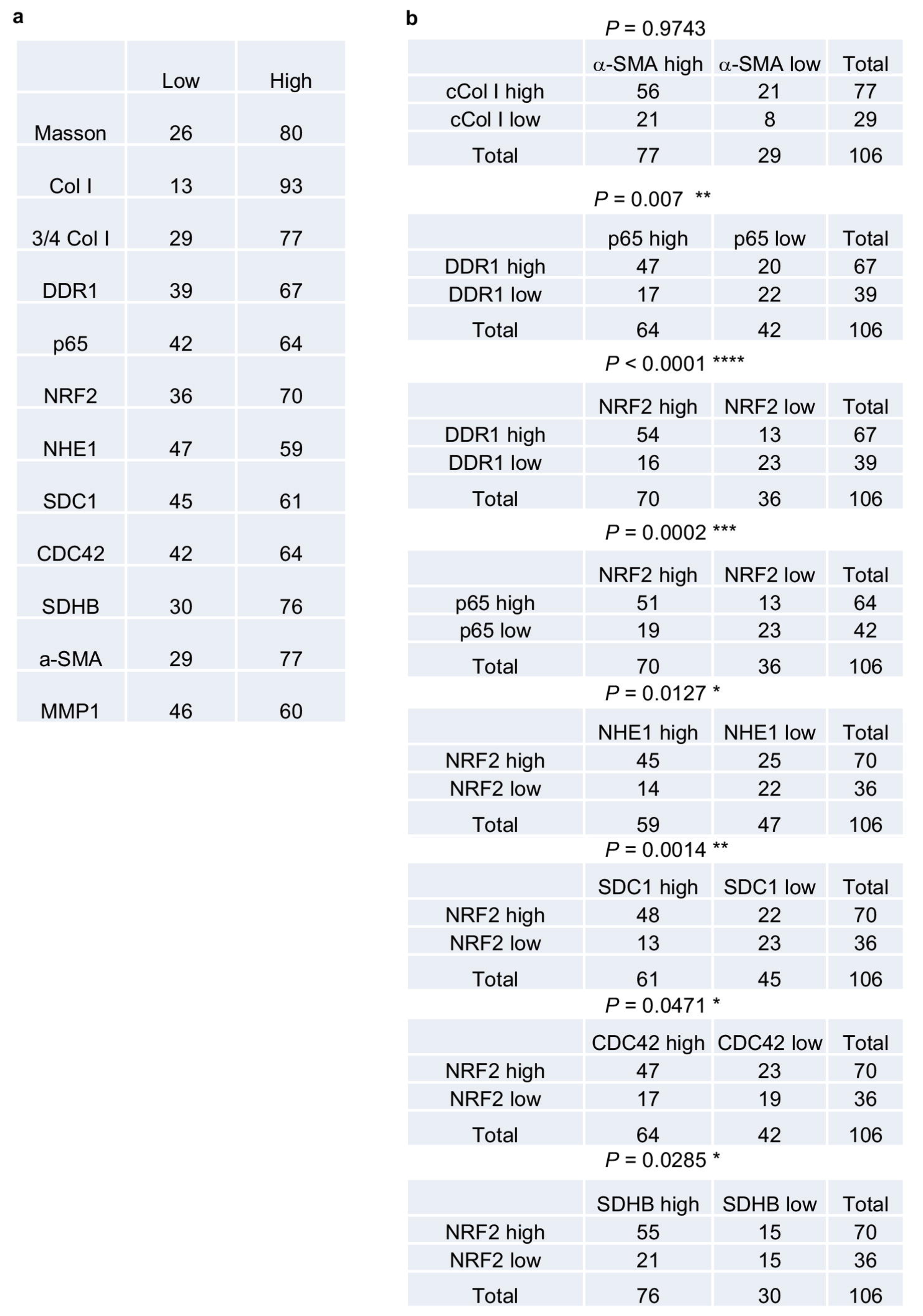
Correlation between Col I-DDR1-NRF2 signaling proteins in human PDAC specimens. **a**, Table depicting numbers and percentages of human PDAC tissues (n=106) positive for the indicated proteins (arbitrarily indicated as low and high). **b**, Correlation between the indicated proteins in the human PDAC tissue array was analyzed by a Chi-square test. *p < 0.05, **p < 0.01, ***p < 0.001, ****p < 0.0001. cCol I (3/4 Col I).

**Extended Data Fig. 7:**
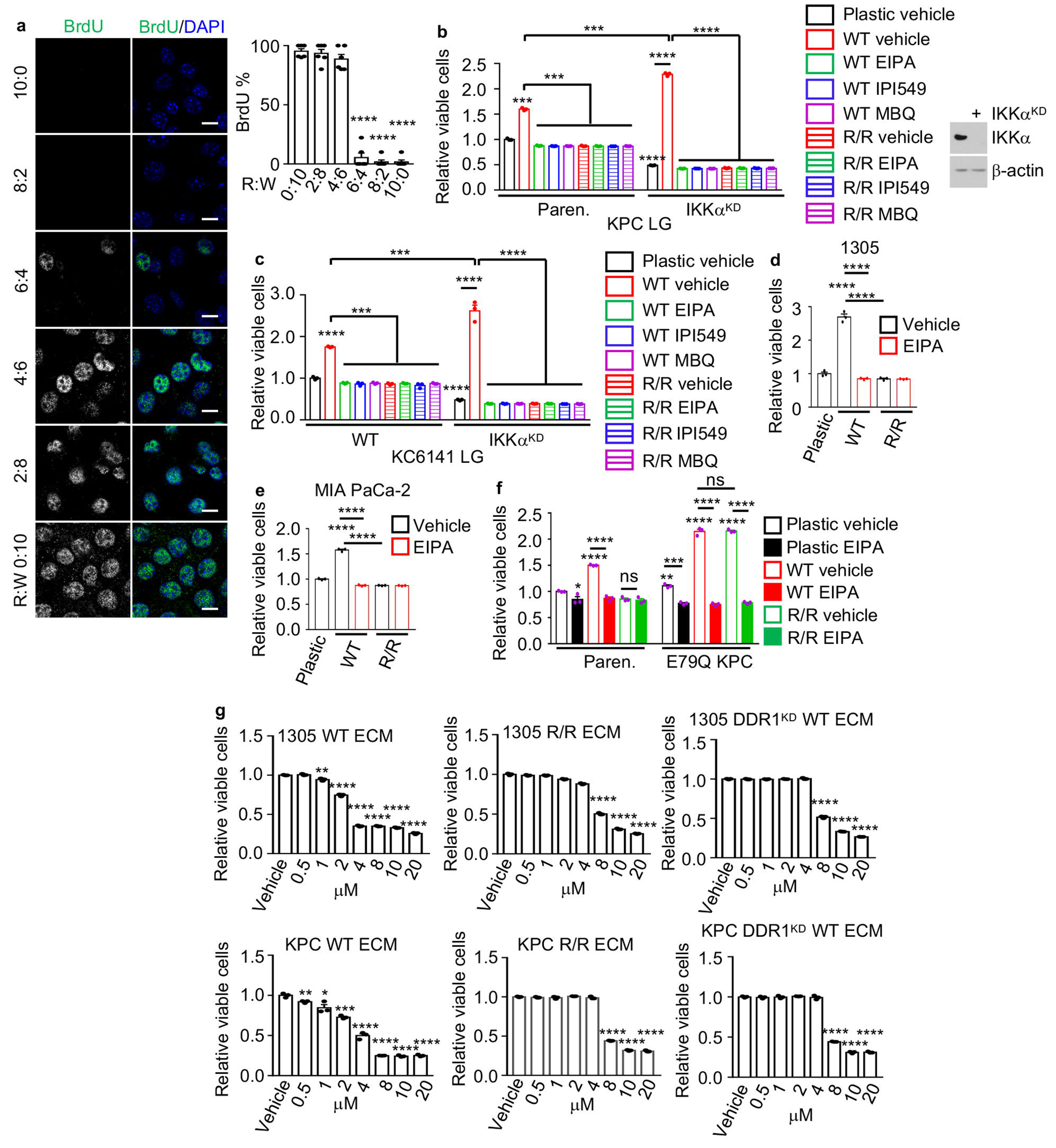
Effect of MP and mitochondria on Col I-controlled PDAC cell growth. **a**, Bromodeoxyuridine (BrdU) incorporation into KPC cells plated on ECM produced by mixtures of R/R and WT fibroblasts (R:W) and incubated in LG plus 0.5 mg/mL BrdU for 24 h. Scale bars, 10 μm. **b**, **c**, Paren. and IKKα^KD^ KPC (**b**) or KC (**c**) cells were plated on plastic, WT or R/R ECM and incubated in LG -/+ EIPA, IPI549 or MBQ. Total viable cells were measured after 3 days, and data are presented relative to untreated Paren. cells plated on plastic. IB analysis of IKKα KD efficiency in KPC cells is shown on the right (**b**). **d**, **e**, 1305 (**d**) or MIA PaCa-2 (**e**) cells were plated as above and incubated in LG -/+ EIPA. Total viable cells were measured as above. **f**, Paren. and E79Q KPC cells were plated and treated as above. Total viable cells were measured and presented as above. **g**, Paren. or DDR1^KD^ 1305 or KPC cells were plated on WT or R/R ECM and incubated in LG medium -/+ the indicated tigecycline concentrations for 24 h. Total viable cells were measured as above, and data are presented relative to the untreated cells. Results in (**a**) (n=6 fields), and (**b**-**g**) (n=3 independent experiments) are mean ± SEM. Statistical significance was determined by a two-tailed t test. *p < 0.05, **p < 0.01, ***p < 0.001, ****p < 0.0001.

**Extended Data Fig. 8:**
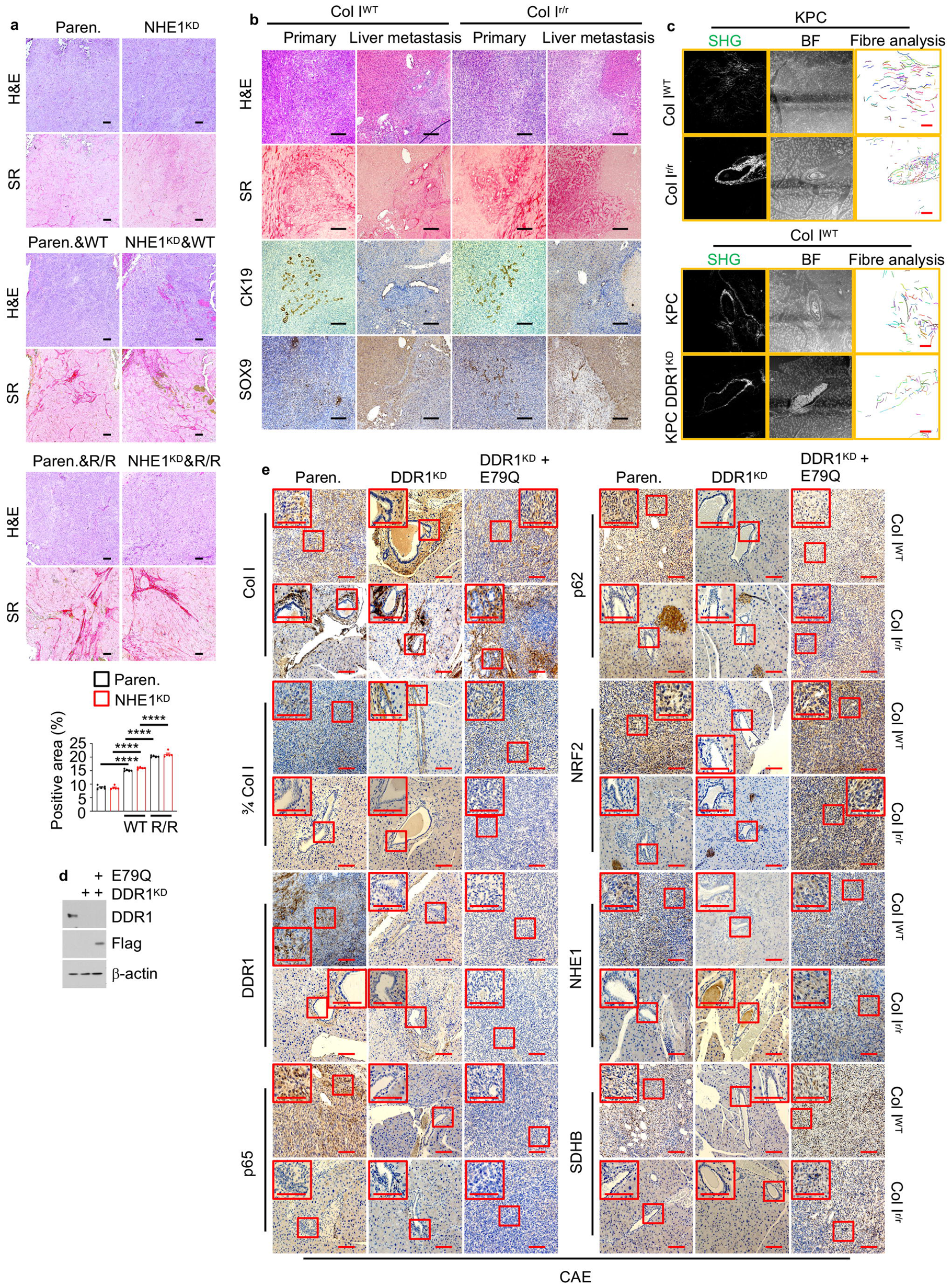
cCol I upregulates MP-related proteins and SDHB via the DDR1-NRF2 axis. **a**, H&E and SR staining of sectioned s.c. tumors formed by Paren. or NHE1^KD^ MIA-PaCa-2 cells transplanted -/+ WT or R/R fibroblasts into nude mice. Quantification of the SR positive area is shown below. Mean ± SEM (n=5). ****p < 0.0001. Scale bars, 100 µm. **b**, H&E, SR and IHC of cytokeratin 19 (CK19) and SOX9 in pancreatic and liver sections from CAE-pretreated Col I^WT^ and Col I^r/r^ mice 4 wk after orthotopic transplantation of KPC NRF2^E79Q^ cells. Scale bars, 100 µm. **c**, Tumors formed by orthotopically transplanted Paren. or DDR1^KD^ KPC cells in Col I^WT^ and Col I^r/r^ mice were analyzed by second harmonic generation (SHG) and collagen fiber individualization. BF-bright field. Scale bars, 100 µm. **d**, IB analysis of DDR1 and Flag-tagged NRF2^E79Q^ in Paren., DDR1^KD^, or NRF2^E79Q^/DDR1^KD^ KPC cells plated on WT ECM in LG medium for 24 h. **e**, IHC of pancreatic sections of CAE pretreated-Col I^WT^ or Col I^r/r^ mice 4 wk after orthotopic transplantation of above cells. Boxed areas were further magnified. Scale bars, 100 µm.

## References

1. Makohon-Moore, A. & Iacobuzio-Donahue, C. A. Pancreatic cancer biology and genetics from an evolutionary perspective. Nat. Rev. Cancer 16, 553–565 (2016).

2. Grzesiak, J. J. & Bouvet, M. The alpha2beta1 integrin mediates the malignant phenotype on type I collagen in pancreatic cancer cell lines. Br. J. Cancer 94, 1311–1319 (2006).

3. Beatty, G. L. et al. CD40 agonists alter tumor stroma and show efficacy against pancreatic carcinoma in mice and humans. Science 331, 1612–1616 (2011).

4. Laklai, H. et al. Genotype tunes pancreatic ductal adenocarcinoma tissue tension to induce matricellular fibrosis and tumor progression. Nat. Med. 22, 497–505 (2016).

5. Incio, J. et al. Obesity-Induced Inflammation and Desmoplasia Promote Pancreatic Cancer Progression and Resistance to Chemotherapy. Cancer Discov. 6, 852–869 (2016).

6. Bissell, M. J. & Hines, W. C. Why don’t we get more cancer? A proposed role of the microenvironment in restraining cancer progression. Nat. Med. 17, 320–329 (2011).

7. Bhattacharjee, S. et al. Tumor restriction by type I collagen opposes tumor-promoting effects of cancer-associated fibroblasts. J. Clin. Invest. 131, 146987 (2021).

8. Rhim, A. D. et al. Stromal elements act to restrain, rather than support, pancreatic ductal adenocarcinoma. Cancer Cell 25, 735–747 (2014).

9. Erkan, M. et al. The activated stroma index is a novel and independent prognostic marker in pancreatic ductal adenocarcinoma. Clin. Gastroenterol. Hepatol. 6, 1155–1161 (2008).

10. Armstrong, T. et al. Type I collagen promotes the malignant phenotype of pancreatic ductal adenocarcinoma. Clin. Cancer Res. 10, 7427–7437 (2004).

11. Ramanathan, R. K. et al. Phase IB/II Randomized Study of FOLFIRINOX Plus Pegylated Recombinant Human Hyaluronidase Versus FOLFIRINOX Alone in Patients With Metastatic Pancreatic Adenocarcinoma: SWOG S1313. J. Clin. Oncol. 37, 1062–1069 (2019).

12. Catenacci, D. V. T. et al. Final analysis of a phase IB/randomized phase II study of gemcitabine (G) plus placebo (P) or vismodegib (V), a hedgehog (Hh) pathway inhibitor, in patients (pts) with metastatic pancreatic cancer (PC): A University of Chicago phase II consortium study. J. Clin. Oncol. 31, 4012–4012 (2013).

13. Wu, H. et al. Generation of collagenase-resistant collagen by site-directed mutagenesis of murine pro alpha 1(I) collagen gene. Proc. Natl. Acad. Sci. U. S. A. 87, 5888–5892 (1990).

14. Zhang, D. Y. & Friedman, S. L. Fibrosis-dependent mechanisms of hepatocarcinogenesis. Hepatol. 56, 769–775 (2012).

15. Shalapour, S. et al. Inflammation-induced IgA+ cells dismantle anti-liver cancer immunity. Nature 551, 340–345 (2017).

16. Baglieri, J. et al. Nondegradable Collagen Increases Liver Fibrosis but Not Hepatocellular Carcinoma in Mice. Am. J. Pathol. 191, 1564–1579 (2021).

17. Su, H. et al. Cancer cells escape autophagy inhibition via NRF2-induced macropinocytosis. Cancer Cell (2021) doi:10.1016/j.ccell.2021.02.016.

18. Criscuolo, D., Avolio, R., Matassa, D. S. & Esposito, F. Targeting Mitochondrial Protein Expression as a Future Approach for Cancer Therapy. Front. Oncol. 11, 797265 (2021).

19. Boraschi-Diaz, I., Wang, J., Mort, J. S. & Komarova, S. V. Collagen Type I as a Ligand for Receptor-Mediated Signaling. Front. Phys. 5, 12 (2017).

20. Vogel, W., Gish, G. D., Alves, F. & Pawson, T. The discoidin domain receptor tyrosine kinases are activated by collagen. Mol. Cell 1, 13–23 (1997).

21. Das, S. et al. Discoidin domain receptor 1 receptor tyrosine kinase induces cyclooxygenase-2 and promotes chemoresistance through nuclear factor-kappaB pathway activation. Cancer Res. 66, 8123–8130 (2006).

22. Zhong, Z. et al. NF-κB Restricts Inflammasome Activation via Elimination of Damaged Mitochondria. Cell 164, 896–910 (2016).

23. Juskaite, V., Corcoran, D. S. & Leitinger, B. Collagen induces activation of DDR1 through lateral dimer association and phosphorylation between dimers. eLife 6, e25716 (2017).

24. Yang, S. H. et al. Discoidin domain receptor 1 is associated with poor prognosis of non-small cell lung carcinomas. Oncol. Rep. 24, 311–319 (2010).

25. Turashvili, G. et al. Novel markers for differentiation of lobular and ductal invasive breast carcinomas by laser microdissection and microarray analysis. BMC Cancer 7, 55 (2007).

26. Barker, K. T. et al. Expression patterns of the novel receptor-like tyrosine kinase, DDR, in human breast tumours. Oncogene 10, 569–575 (1995).

27. Heinzelmann-Schwarz, V. A. et al. Overexpression of the cell adhesion molecules DDR1, Claudin 3, and Ep-CAM in metaplastic ovarian epithelium and ovarian cancer. Clin. Cancer Res. 10, 4427–4436 (2004).

28. Sun, X. et al. Tumour DDR1 promotes collagen fibre alignment to instigate immune exclusion. Nature 599, 673–678 (2021).

29. Di Martino, J. S. et al. A tumor-derived type III collagen-rich ECM niche regulates tumor cell dormancy. *Nat*. Cancer (2021) doi:10.1038/s43018-021-00291-9.

30. Picca, A. & Lezza, A. M. S. Regulation of mitochondrial biogenesis through TFAM-mitochondrial DNA interactions: Useful insights from aging and calorie restriction studies. Mitochondrion 25, 67–75 (2015).

31. Scarpulla, R. C., Vega, R. B. & Kelly, D. P. Transcriptional integration of mitochondrial biogenesis. Trends Endocrinol. Metab. 23, 459–466 (2012).

32. Aguilera, K. Y. et al. Inhibition of Discoidin Domain Receptor 1 Reduces Collagen-mediated Tumorigenicity in Pancreatic Ductal Adenocarcinoma. Mol. Cancer Ther. 16, 2473–2485 (2017).

33. Vogel, W. Discoidin domain receptors: structural relations and functional implications. FASEB J. 13 Suppl, S77-82 (1999).

34. Whatcott, C. J. et al. Desmoplasia in Primary Tumors and Metastatic Lesions of Pancreatic Cancer. Clin. Cancer Res. 21, 3561–3568 (2015).

35. Hwang, R. F. et al. Cancer-associated stromal fibroblasts promote pancreatic tumor progression. Cancer Res. 68, 918–926 (2008).

36. Mahajan, U. M. et al. Immune Cell and Stromal Signature Associated With Progression-Free Survival of Patients With Resected Pancreatic Ductal Adenocarcinoma. Gastroenterology 155, 1625–1639.e2 (2018).

37. Bachem, M. G. et al. Pancreatic carcinoma cells induce fibrosis by stimulating proliferation and matrix synthesis of stellate cells. Gastroenterology 128, 907–921 (2005).

38. Chen, Y. et al. Type I collagen deletion in αSMA+ myofibroblasts augments immune suppression and accelerates progression of pancreatic cancer. Cancer Cell (2021) doi:10.1016/j.ccell.2021.02.007.

39. Affo, S. et al. Promotion of cholangiocarcinoma growth by diverse cancer-associated fibroblast subpopulations. Cancer Cell (2021) doi:10.1016/j.ccell.2021.03.012.

40. Zhang, T., Shen, S., Qu, J. & Ghaemmaghami, S. Global Analysis of Cellular Protein Flux Quantifies the Selectivity of Basal Autophagy. Cell Rep. 14, 2426–2439 (2016).

41. Liu, X. et al. A targeted mutation at the known collagenase cleavage site in mouse type I collagen impairs tissue remodeling. J. Cell Biol. 130, 227–237 (1995).

42. Soares, K. C. et al. A preclinical murine model of hepatic metastases. J. Vis. Exp. (2014) doi:10.3791/51677.

43. Su, H. et al. VPS34 Acetylation Controls Its Lipid Kinase Activity and the Initiation of Canonical and Non-canonical Autophagy. Mol. Cell 67, 907–921.e7 (2017).

44. Lee, S.-W., Alas, B. & Commisso, C. Detection and Quantification of Macropinosomes in Pancreatic Tumors. Methods Mol. Biol. 171–181 (2019).

45. Monaghan, M. G., Kroll, S., Brucker, S. Y. & Schenke-Layland, K. Enabling Multiphoton and Second Harmonic Generation Imaging in Paraffin-Embedded and Histologically Stained Sections. Tissue Eng. Part C Methods 22, 517–523 (2016).

46. Lee, J. J. et al. Elucidation of Tumor-Stromal Heterogeneity and the Ligand-Receptor Interactome by Single-Cell Transcriptomics in Real-world Pancreatic Cancer Biopsies. Clin. Cancer Res. 27, 5912–5921 (2021).

47. Hao, Y. et al. Integrated analysis of multimodal single-cell data. Cell 184, 3573–3587.e29 (2021).

48. Kumar, A., Mitchener, J., King, Z. A. & Metallo, C. M. Escher-Trace: a web application for pathway-based visualization of stable isotope tracing data. BMC Bioinformatics 21, 297 (2020).

49. Commisso, C., Flinn, R. J. & Bar-Sagi, D. Determining the macropinocytic index of cells through a quantitative image-based assay. Nat. Protoc. 9, 182–192 (2014).

